# Experience-dependent diversification of ripple firing and synaptic inputs in hippocampal CA1

**DOI:** 10.64898/2025.12.26.696628

**Authors:** Junko Ishikawa, Takuto Tomokage, Dai Mitsushima

## Abstract

The hippocampus plays a key role in encoding episodic memory by transforming recent experience into persistent neuronal and synaptic modifications. However, the physiological processes that link real-world experience to coordinated network activity and synaptic reorganization remain incompletely understood. Here, we investigated how distinct types of episodic experience reshape ensemble firing dynamics and synaptic input in the hippocampal CA1 region of freely moving rats.

We identified spontaneous *super bursts*, defined as brief episodes of high-frequency population firing, that emerged preferentially during emotionally salient experiences. These bursts were associated with an increase in ripple firing, defined as short-duration, high-frequency multi-unit spike activity occurring in association with sharp-wave ripples. Analysis of ripple firing patterns revealed experience-dependent diversification, reflected by increased information entropy after episodic experience. Ex vivo whole-cell patch-clamp recordings further demonstrated that miniature excitatory and inhibitory synaptic currents in CA1 pyramidal neurons underwent experience-specific reorganization.

Together, these findings support a coordinated cascade in which episodic experience induces population-level ensemble activity, followed by diversification of ripple firing patterns and reorganization of excitatory and inhibitory synaptic inputs in hippocampal CA1. This coordination defines a population-level signature associated with experience-dependent encoding across hippocampal circuits.

**Key points:**

- The hippocampus is essential for episodic memory, but how real-life experiences change brain activity and synaptic connections remains unclear.
- In freely moving rats, emotionally salient experiences triggered brief bursts of high-frequency firing involving many neurons in the hippocampal CA1 region (“super bursts”).
- After these experiences, short high-frequency firing events linked to memory processing (“ripple firing”) became more diverse in their timing and shape.
- Recordings from individual neurons showed that both excitatory and inhibitory synaptic inputs were reorganized in an experience-specific manner.
- These results suggest that episodic experience is encoded through coordinated changes in population activity, ripple firing patterns, and synaptic inputs in hippocampal CA1.

## INTRODUCTION

Every day, animals encounter events that combine elements of “when, where, and what.” The hippocampus has long been recognized as a critical structure involved in episodic memory processing (1), supporting the representation of spatiotemporal information (2,3) and specific experiential contexts (4). Within the dorsal hippocampus, CA1 neurons are known to exhibit experience-dependent activity related to spatial context and novelty (5), and transient disruption of CA1 function impairs memory-guided behavior (6). Consistent with this, learning to avoid aversive environments requires synaptic plasticity in dorsal CA1 neurons (7,8). CA1 neurons also contribute to object recognition (9) and social representations, such as encoding information about conspecific location during social interaction (10). Despite these advances, how distinct episodic experiences are differentially reflected in hippocampal neuronal activity and retained over time remains incompletely understood.

CA1 neurons frequently engage in coordinated firing with neighboring neurons, and the extent of synchronization increases with behavioral demands (11). Because spatiotemporal firing patterns across neuronal populations can convey information (12,13), hippocampal activity has been studied extensively using large-scale electrophysiological approaches (14). In particular, sharp-wave ripple (SPW-R)–associated activity has been proposed to play an important role in memory-related processing (15). Experimental disruption of SPW-Rs impairs memory acquisition and retrieval, whereas successful learning enhances ripple duration and neuronal participation (16–18). However, capturing coordinated spike activity among neighboring neurons during ripple-associated events with sufficient temporal resolution remains technically challenging. Moreover, spike waveform properties can vary over time within individual neurons, especially during burst firing (19), and learning-related dendritic plateau potentials can generate complex firing patterns (20). These considerations motivate approaches that focus on population-level firing dynamics rather than strict single-unit isolation when examining experience-dependent hippocampal activity in freely moving animals.

Emotional states such as fear, reward, and stress strongly influence memory formation (21–24), and these effects are mediated in part by neuromodulatory transmission in the hippocampus. Emotional arousal enhances learning through noradrenergic activation of dorsal CA1 neurons, promoting synaptic incorporation of GluA1-containing AMPA receptors (25). In addition, locus coeruleus neurons expressing tyrosine hydroxylase may facilitate memory by co-releasing dopamine in the hippocampus (26). Nevertheless, direct evidence linking emotional experience to dynamic changes in CA1 population firing patterns in freely moving animals remains limited. Furthermore, although a causal relationship between synaptic plasticity and learning has been well established (7,8,27), spontaneous high-frequency population firing associated with learning has been less systematically characterized.

In addition to excitatory plasticity, inhibitory circuit maturation and plasticity are essential for hippocampal learning (28,29). We previously identified cell- and pathway-specific synaptic modifications that correlate with acquired behavioral performance (30,31). Notably, synaptic plasticity induced during the initial phase of experience rapidly generates diversity in both excitatory and inhibitory postsynaptic currents within minutes, and these changes persist for extended periods (32). Such rapid synaptic diversification may provide a substrate for differentiating distinct episodic experiences (33).

In the present study, we combined multi-unit recordings of CA1 neuronal activity in freely moving rats with *ex vivo* patch-clamp analyses to examine how episodic experiences with different emotional content are associated with coordinated population firing and synaptic reorganization. By analyzing experience-dependent changes in ensemble firing dynamics and synaptic inputs, we sought to provide an integrated physiological framework linking hippocampal network activity with cellular-level plasticity.

## RESULTS

### Behavioral evidence of experience-dependent memory

To model episodic experiences in rodents, adult male rats were exposed for 10 min to one of four experimental episodes (Fig. 1A): restraint stress, social interaction with a female or a male, or observation of a novel object (Fig. 1B). To assess experience-dependent memory, animals were re-exposed to the same episode on the following day, and their behavior was evaluated (34,35).

**Figure 1.**
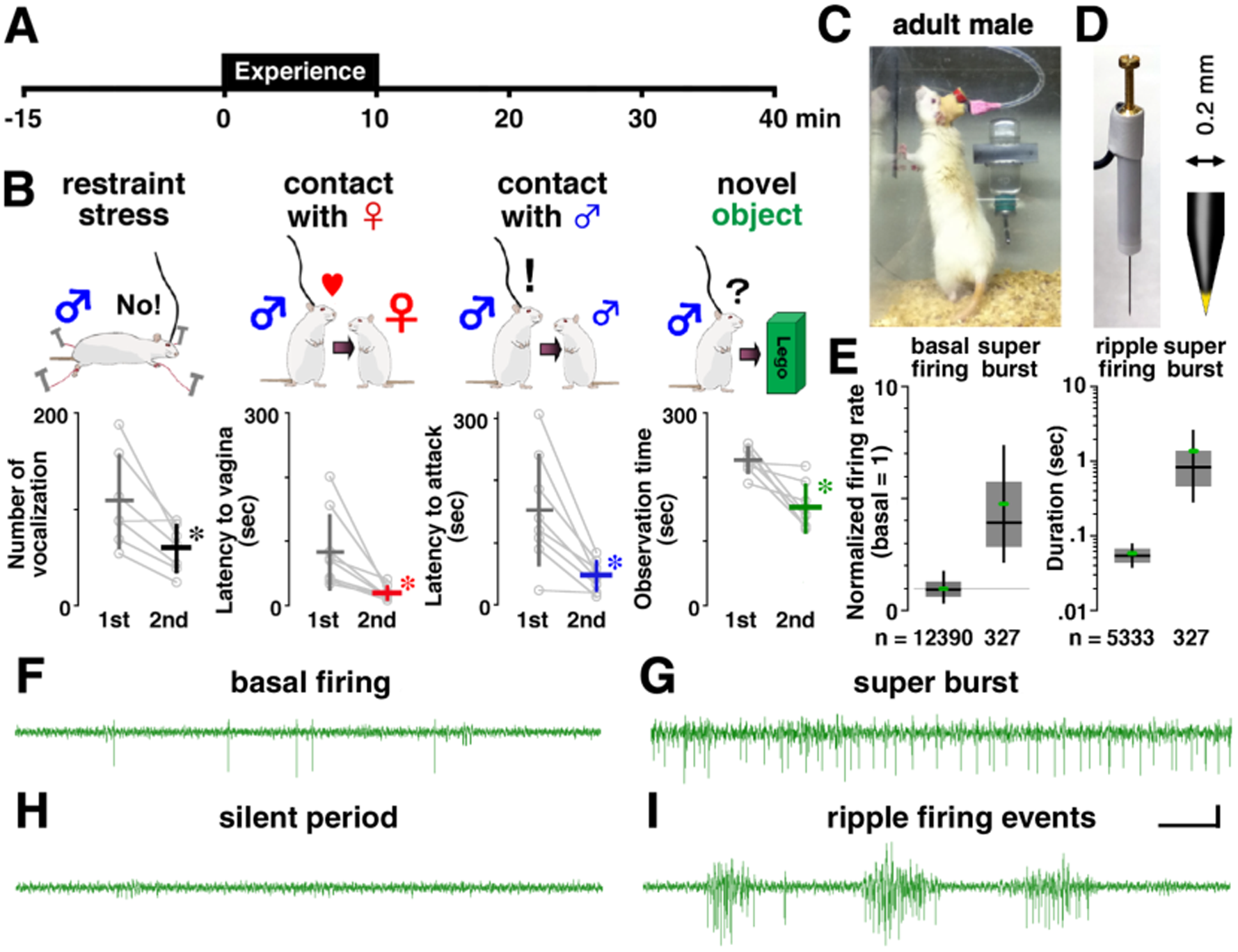
Experimental design for episodic memory and classification of CA1 multi-unit firing patterns. **(A)** Timeline of recording sessions and 10-min episodic exposures. **(B)** Schematic illustrations of the four episodic experiences. Memory acquisition was assessed on the following day (2nd exposure). Data are shown as mean ± SD; gray circles represent individual rats, with lines indicating within-subject changes. *P < 0.05 vs. 1st exposure. **(C)** Photograph of a recorded animal. **(D)** Movable recording electrode with an enlarged tip. **(E)** Super bursts were extracted as events exceeding 3 SD of basal firing rate are shown as normalized firing rate (basal = 1) (left). Ripple firing events were distinguished from super bursts based on event duration (right). Boxes indicate the interquartile range with the median shown as a horizontal line; whiskers indicate the minimum and maximum values excluding outliers. Green dots indicate mean values. Sample size (number of events) is shown below each bar. **(F–I)** Examples of multiple-unit activity from hippocampal CA1 at 25 kHz and filtered between 300–10,000 Hz. All traces were recorded from an electrode implanted in the same animal. **(F)** Basal firing, **(G)** super burst, **(H)** silent period, and **(I)** ripple firing events (three examples) within the same recording. Each event displayed a distinct waveform pattern. Scale bar = 50 ms. Note: Spike classification using the Spike2 algorithm is unreliable during super bursts and ripple firing events (see Fig. S1).

Rats subjected to restraint stress exhibited fewer vocalizations during the second exposure (t_6_ = 3.476, P = 0.0129). Similarly, rats exposed to a female, male, or novel object showed reduced latency to vaginal inspection (t_8_ = 3.492, P = 0.0082), attack (t_7_ = 4.192, P = 0.0041), or object observation time (t_9_ = 2.901, P = 0.0176), respectively, during the second encounter (36). These behavioral changes were consistent with experience-dependent memory across the different episodic conditions.

### Multiple-unit activity and classification of firing patterns

To examine experience-dependent neuronal activity, we recorded multiple-unit firing from the CA1 region before (15 min), during (10 min), and after (30 min) each episode using chronically implanted electrodes capable of detecting activity from neighboring neurons (Figs. 1C, D). The number of recordings was as follows: no-experience control (N = 7), restraint stress (N = 9), contact with a female (N = 11), contact with a male (N = 11), and novel object (N = 9).

Because individual neurons could not be reliably isolated from the multi-unit signals (Fig. S1A-C), all quantitative analyses were performed at the population level. Spike waveform analysis was conducted only to confirm recording location within the pyramidal cell layer. Based on spike half-width during basal firing, 284 well-separated waveforms were classified as putative pyramidal neurons (>0.7 ms; n = 220), interneurons (<0.4 ms; n = 38), or unclassified units (0.4–0.7 ms; n = 26), supporting electrode placement in CA1 (Fig. S1D). These classifications were not used for subsequent quantitative analyses. Neighboring CA1 neurons exhibited distinct firing patterns, particularly following the onset of episodic experience. Based on predefined criteria, we extracted two types of population firing events: super burst events and ripple firing events (Fig. 1E).

Before episodic exposure, CA1 activity consisted primarily of sporadic firing, with occasional ripple firing events in the home cage (Fig. 1F). Baseline firing rates were calculated from 100–300 s of low-noise data prior to experience. Super burst events were defined as spontaneous high-frequency population firing with rates exceeding three standard deviations (SD) above baseline (n = 100–300 events per recording; Figs. 1G, 2A,B).

**Figure 2.**
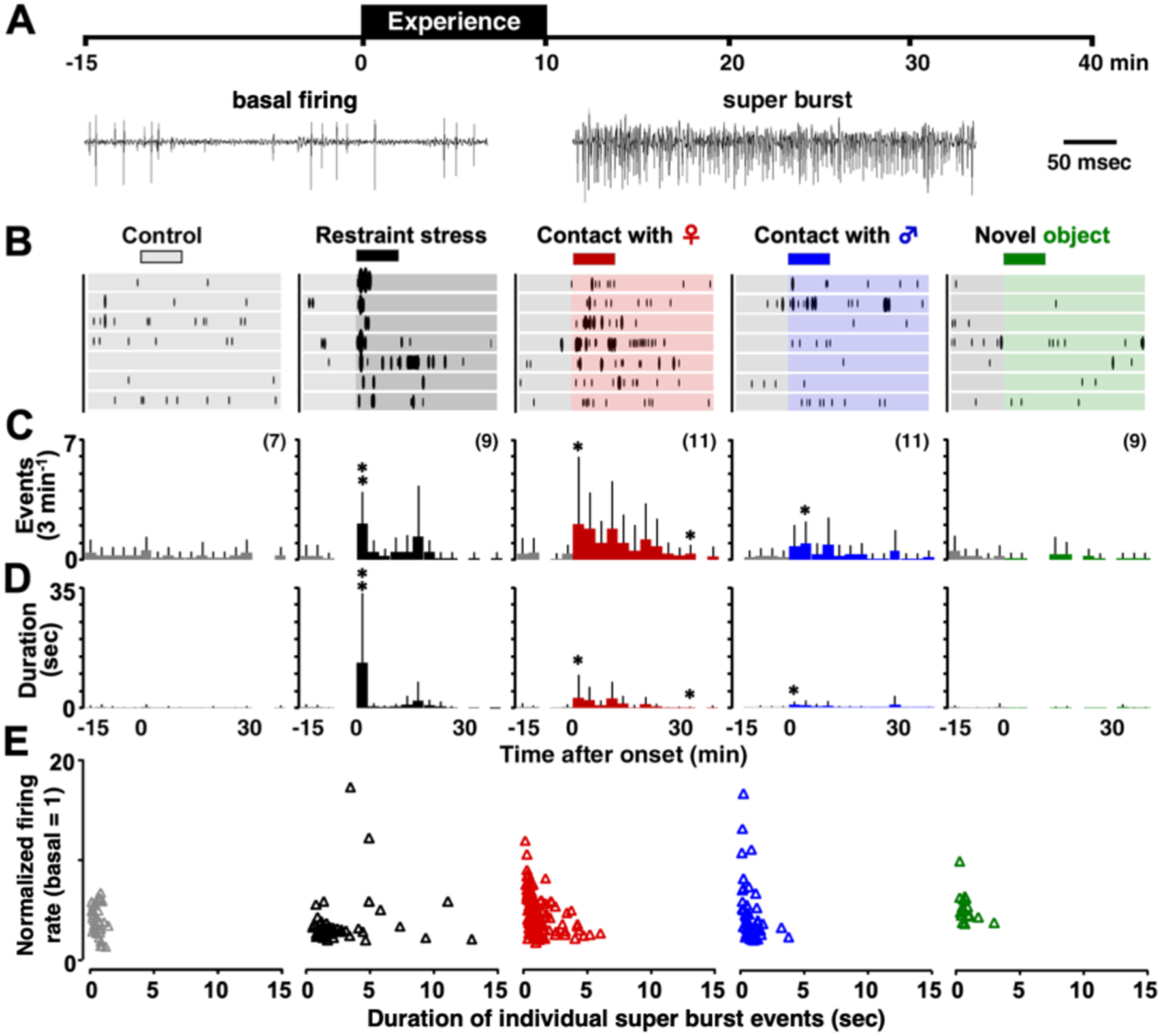
Experience-dependent induction of high-frequency CA1 multi-unit firing (super bursts) **(A)** Representative traces showing basal firing and an episode-induced super burst recorded from CA1 neurons. **(B)** Time course of super burst occurrences before, during, and after episodic experiences in individual animals. Horizontal black bars indicate the 10-min episode window. **(C–D)** Quantification of the occurrence (C) and total duration (D) of super bursts in 3-min bins across time. **(E)** Scatter plots of individual super burst events, showing episode-specific differences in event duration (x-axis) and normalized firing rate (y-axis; basal = 1). Each triangle represents a single super burst event. Multivariate analysis of these two event-level features revealed distinct episode-dependent patterns (Table 1, MANOVA). Data are shown as mean ± SD. The number of recordings in each group is shown in parentheses. *P < 0.05, **P < 0.01 vs. pre-experience baseline.

**Table 1.** Experience-specific super burst patterns revealed by multivariate analysis (related to Fig. 2).

| Comparison | Incidence–Duration | Frequency–Duration |
| --- | --- | --- |
| Overall | $F_{8, 1304} = 9.761, P < 0.001^{**}$ | $F_{8, 558} = 11.535, P < 0.001^{**}$ |
| Restraint vs Female | $F_{2, 277} = 14.066, P < 0.001^{**}$ | $F_{2, 177} = 24.009, P < 0.001^{**}$ |
| Restraint vs Male | $F_{2, 277} = 12.948, P < 0.001^{**}$ | $F_{2, 107} = 20.873, P < 0.001^{**}$ |
| Restraint vs Object | $F_{2, 249} = 8.367, P < 0.001^{**}$ | $F_{2, 66} = 19.088, P < 0.001^{**}$ |
| Restraint vs Control | $F_{2, 221} = 9.517, P < 0.001^{**}$ | $F_{2, 82} = 33.336, P < 0.001^{**}$ |
| Female vs Male | $F_{2, 305} = 4.793, P < 0.001^{**}$ | $F_{2, 181} = 6.037, P = 0.003^{**}$ |
| Female vs Object | $F_{2, 277} = 13.075, P < 0.001^{**}$ | $F_{2, 140} = 1.589, P = 0.208$ |
| Female vs Control | $F_{2, 249} = 7.004, P < 0.001^{**}$ | $F_{2, 156} = 4.695, P = 0.011^{*}$ |
| Male vs Object | $F_{2, 277} = 5.160, P = 0.006^{**}$ | $F_{2, 70} = 1.912, P = 0.155$ |
| Male vs Control | $F_{2, 249} = 1.248, P = 0.289$ | $F_{2, 86} = 2.806, P = 0.066$ |
| Object vs Control | $F_{2, 221} = 1.916, P = 0.150$ | $F_{2, 45} = 0.445, P = 0.644$ |

MANOVA and post hoc analyses comparing integrated super burst features (incidence–duration and frequency–duration relationships) across episodic experiences. Significant differences indicate experience-specific modulation of ensemble-level high-frequency population firing.

Ripple firing events were detected in association with SPW-Rs (150–300 Hz). We analyzed the 300–10 kHz band to characterize clustered spike activity occurring during these oscillations. Ripple firing events were short-duration (56.3 ± 16.5 ms, mean ± SD; n = 5333), high-frequency population firing events with a signal-to-noise ratio ≥6:1 (Fig. 1I), typically separated by silent periods with no firing (Fig. 1H). To provide an intuitive overview of CA1 population firing patterns, representative examples of multi-unit activity before and after episodic experience are shown in Movie S1 and Movie S2, respectively.

### Super burst activity shows episode-type specific differences

We examined changes in the occurrence and duration of super burst events across time and episodic conditions (Figs. 2C,D). Two-way ANOVA with experience as the between-group factor and time as the within-group factor revealed significant effects of episode (duration: *F*_4, 546_ = 3.145, *P* = 0.024), time (events: *F*_13, 546_ = 4.937, *P* < 0.0001; duration: *F*_13, 546_ = 7.288, *P* < 0.0001), and their interaction (events: *F*_52, 546_ = 1.567, *P* = 0.009; duration: *F*_52, 546_ = 3.095, *P* < 0.0001).

Post hoc analyses showed that both occurrence and duration of super burst events increased following restraint stress, contact with a female, and contact with a male, whereas no significant changes were observed after exposure to a novel object or in no-experience controls. Multivariate analysis further revealed episode-type specific differences in super burst occurrence and duration across time (two-way MANOVA: *F*_8,1304_ = 9.761, *P* < 0.0001; Table 1).

Analysis of individual super burst events demonstrated that the relationship between duration and relative firing rate differed between episodic conditions (Fig. 2E). One-way MANOVA revealed episode-dependent differences in integrated super burst features (*F*_8, 558_ = 11.535, *P* < 0.0001).

### Episodic experience increases silent periods across conditions

Silent periods, defined as intervals without firing exceeding baseline variability, were examined across episodes (Fig. 3A,B). Two-way ANOVA revealed a significant effect of time (*F*_2,84_ = 18.167, *P* < 0.0001), but neither the main effect of experience nor the interaction was significant.

**Figure 3.**
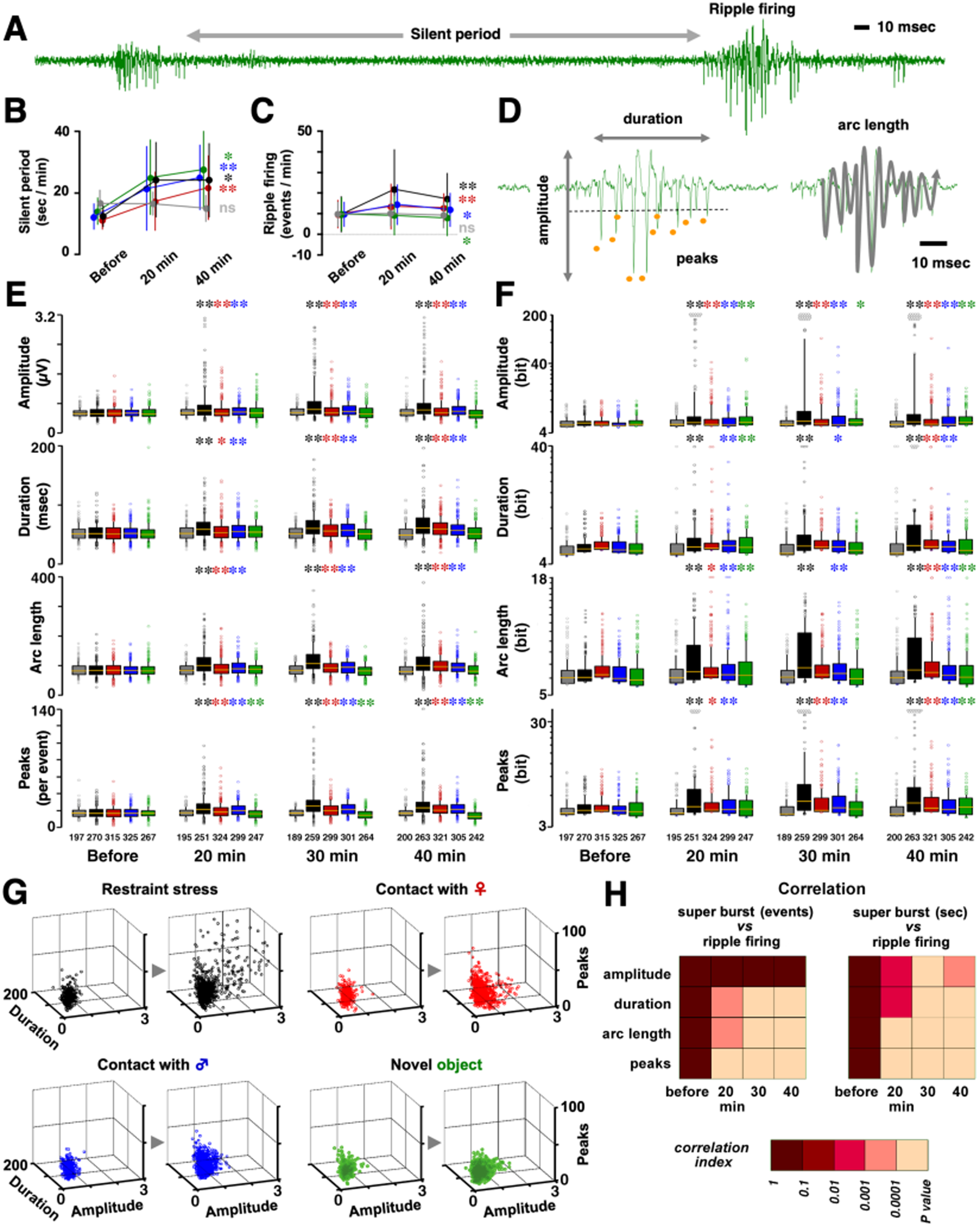
Experience-dependent diversification of ripple firing events in CA1 neurons. **(A)** Representative trace showing a silent period flanked by two ripple firing events. **(B–C)** Episodic experience increased the total duration of silent periods (**B**) and the incidence of ripple firing events (**C**). Data are shown as mean ± SD. (**D**) Four quantitative features extracted from individual ripple firing events: amplitude, duration, arc length, and number of negative peaks. (**E**) Temporal dynamics of ripple firing features across conditions, showing episode-dependent modulation. Box plots indicate the interquartile range (50%), median, and minimum–maximum values excluding outliers. Sample size (number of ripple firing events) is shown below each bar. (**F**) Information entropy per ripple firing event, calculated from the distribution of waveform features, increased following episodic experience. (**G**) Multidimensional analysis integrating the four ripple firing features revealed episode-type-specific clustering patterns (Table 2, MANOVA; three representative dimensions are shown). (**H**) Correlations between super burst parameters and ripple firing features before (left) and after (right) episodic experience. Significant correlations emerged only after experience. *P < 0.05, **P < 0.01 vs. pre-experience.

**Table 2.** Experience-specific diversification of ripple firing features revealed by multivariate analysis (related to Fig. 3).

| Comparison | Features | Self-entropy |
| --- | --- | --- |
| Overall | $F_{12, 11993} = 48.996, P < 0.001^{**}$ | $F_{16, 16223} = 35.748, P < 0.001^{**}$ |
| Restraint vs Female | $F_{4, 2291} = 30.810, P < 0.001^{**}$ | $F_{4, 2291} = 33.374, P < 0.001^{**}$ |
| Restraint vs Male | $F_{4, 2262} = 31.515, P < 0.001^{**}$ | $F_{4, 2262} = 42.701, P < 0.001^{**}$ |
| Restraint vs Object | $F_{4, 2052} = 86.412, P < 0.001^{**}$ | $F_{4, 2052} = 48.154, P < 0.001^{**}$ |
| Restraint vs Control | $F_{4, 1813} = 48.846, P < 0.001^{**}$ | $F_{4, 1813} = 52.735, P < 0.001^{**}$ |
| Female vs Male | $F_{4, 2478} = 3.395, P = 0.009^{**}$ | $F_{4, 2478} = 3.920, P = 0.004^{**}$ |
| Female vs Object | $F_{4, 2268} = 60.959, P < 0.001^{**}$ | $F_{4, 2268} = 12.403, P < 0.001^{**}$ |
| Female vs Control | $F_{4, 2029} = 18.258, P < 0.001^{**}$ | $F_{4, 2029} = 43.749, P < 0.001^{**}$ |
| Male vs Object | $F_{4, 2239} = 94.403, P < 0.001^{**}$ | $F_{4, 2239} = 6.812, P < 0.001^{**}$ |
| Male vs Control | $F_{4, 2000} = 32.546, P < 0.001^{**}$ | $F_{4, 2000} = 31.836, P < 0.001^{**}$ |
| Object vs Control | $F_{4, 1790} = 23.418, P < 0.001^{**}$ | $F_{4, 1790} = 29.895, P < 0.001^{**}$ |

MANOVA and post hoc analyses comparing integrated ripple firing features (amplitude, duration, arc length, and number of negative peaks) across episodic experiences. Detailed results are provided in Tables S5 (integrated features) and S6 (integrated self-entropy). Significant differences indicate experience-specific diversification of ripple firing features.

Within-group analyses showed that silent periods increased following all episodic experiences, including restraint stress, social interactions, and novel object exposure, whereas no significant changes were observed in no-experience controls (Table S1). These results indicate that episodic experience consistently increased silent periods irrespective of episode type.

### Ripple firing dynamics depend on episodic experience

Ripple oscillations are known hippocampal network events associated with high-frequency population firing. To assess experience-dependent changes in ripple firing events, we quantified their occurrence across time (Fig. 3C).

Two-way ANOVA revealed significant effects of time (*F*_2,84_ = 10.649, *P* < 0.0001) and interaction (*F*_8,84_ = 3.422, *P* = 0.002), but not experience alone. Within-group analyses demonstrated increased ripple firing occurrence following restraint stress and social interactions, whereas exposure to a novel object resulted in a decrease. No significant changes were observed in no-experience controls (Table S2).

### Super burst activity is associated with ripple firing but not silent periods

Because depolarization is a prerequisite for synaptic plasticity, we examined whether super burst activity was associated with changes in ripple firing or silent period. Across animals, changes in silent periods were not correlated with the number or total duration of super burst events (Figs. S3A, B). In contrast, both the number and total duration of super burst events were positively correlated with changes in ripple firing occurrence (Figs. S3C, D).

### Experience-dependent diversification of ripple firing features

Individual ripple firing events were characterized by four features (Fig. 3D). Two-way repeated-measures ANOVA revealed significant experience- and time-dependent changes across features (Fig. 3E; Table S3). Most features increased following episodic experience, whereas the number of peaks decreased specifically after exposure to a novel object. These changes persisted for at least 40 min and were absent in no-experience controls. We further quantified the diversity of ripple firing features using Shannon entropy (Fig. 3F). Information entropy per ripple firing event increased significantly following episodic experience and remained elevated for at least 40 min. Two-way repeated-measures ANOVA confirmed significant effects across episodes and time (Table S4). Multivariate analyses integrating the four ripple firing features revealed episode-type specific differences in ripple firing diversification (Fig. 3G; Table 2). Similar results were obtained when analyzing integrated self-entropy values. Notably, experience-induced changes in ripple firing features and entropy were significantly correlated with preceding super burst activity (Fig. 3H), whereas ripple firing features observed before experience showed no such relationship.

Because ripple firing reflects coordinated activity within CA1 local circuits, we next examined whether the experience-dependent diversification observed at the level of ripple firing events was accompanied by corresponding changes in excitatory and inhibitory synaptic inputs onto CA1 pyramidal neurons.

### Episode-type specific diversification of synaptic currents in CA1 neurons

To assess synaptic changes following episodic experience, we performed ex vivo whole-cell recordings from CA1 pyramidal neurons 40 min after exposure. Sequential recordings of mEPSCs and mIPSCs yielded four parameters per neuron: amplitude and frequency for each current type (Fig. 4A).

**Figure 4.**
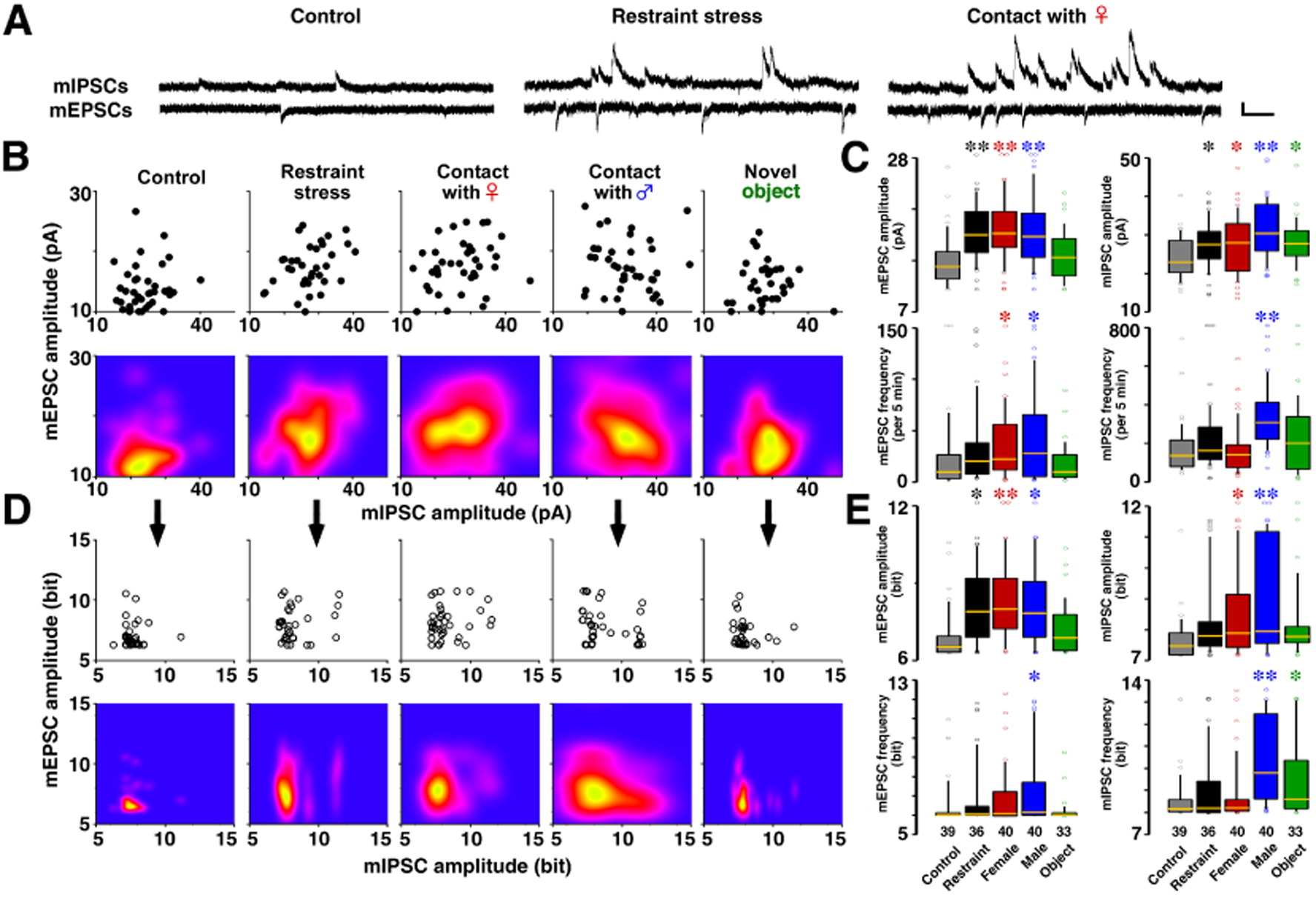
Experience-dependent diversification of excitatory and inhibitory synaptic inputs in CA1 neurons. (**A**) Representative traces of miniature excitatory (mEPSCs, top) and inhibitory (mIPSCs, bottom) postsynaptic currents recorded from the same CA1 pyramidal neuron. (**B**) Two-dimensional representation of synaptic input strength showing the relationship between mEPSC and mIPSC amplitudes (top, scatter plots) and their corresponding density distributions (bottom). Although only two dimensions are shown, multidimensional analysis integrating all four synaptic parameters (amplitude and frequency of mEPSCs and mIPSCs) revealed episode-type-specific synaptic patterns (Table 3, MANOVA). (**C**) Box plots summarizing experience-dependent changes in the four synaptic parameters: amplitude and frequency of mEPSCs and mIPSCs (ANOVA). (**D**) Information entropy per neuron, calculated from the joint distribution of the four synaptic parameters, exhibited experience- and episode-type-specific diversification (Table 3, MANOVA). (**E**) Box plots of individual self-entropy components derived from each synaptic parameter (ANOVA). Boxes indicate the interquartile range with the median shown as a line; whiskers represent minimum to maximum values excluding outliers. Sample size (number of neurons) is shown below each bar. *P < 0.05, **P < 0.01 vs. control.

**Table 3.**
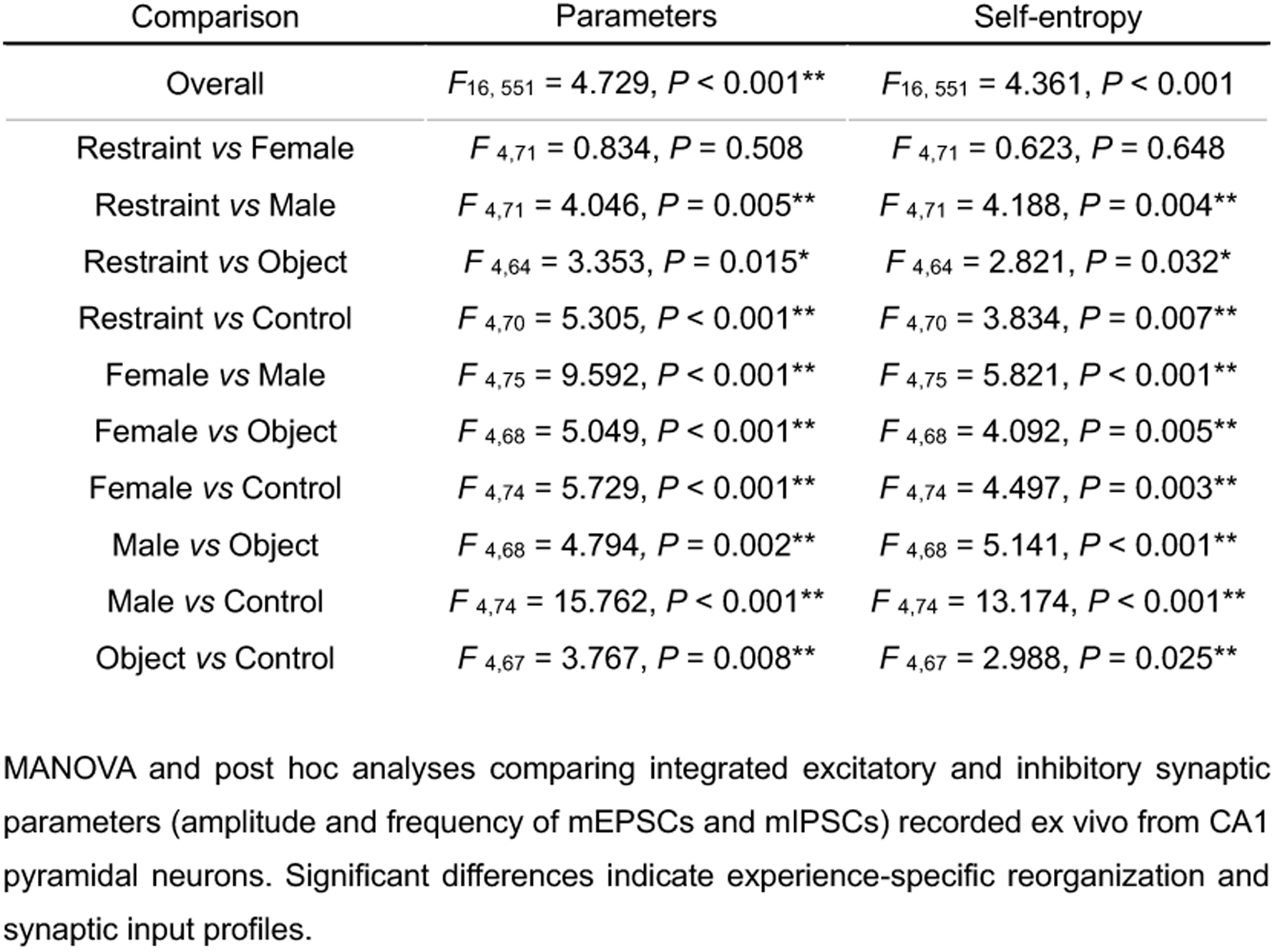
Experience-specific reorganization of excitatory and inhibitory synaptic inputs revealed by multivariate analysis (related to Fig. 4).

Experience diversified the distribution of synaptic parameters compared with no-experience controls (Figs. 4B–D). One-way ANOVA revealed episode-dependent changes across individual parameters (Fig. 4C; Table S7). Multivariate analysis integrating all four parameters demonstrated episode-type specific synaptic reorganization (MANOVA: F₁₆,₅₅₁ = 4.729, P < 0.001; Fig. 4B; Table 3). Shannon entropy analysis further showed that synaptic input diversity increased following episodic experience (Fig. 4E; Table S8). Integrated self-entropy analysis confirmed episode-type specific diversification of excitatory and inhibitory synaptic inputs (Fig. 4D).

## DISCUSSION

Long-term potentiation (LTP) is widely accepted as a synaptic model of learning and memory (37) and is now supported by causal optogenetic evidence (8,23). Despite this progress, a fundamental gap remains between experimentally imposed induction paradigms and natural learning conditions. The endogenous population-level activity patterns that transform experience into persistent synaptic plasticity in behaving animals remain poorly understood, particularly across multiple organizational levels.

In the present study, we focused on spontaneous high-frequency population firing events in hippocampal CA1 neurons that emerged preferentially during emotionally salient experiences. These events, termed *super bursts*, were defined as transient increases in population firing exceeding baseline variability and were examined in relation to subsequent changes in ripple firing patterns and synaptic organization. Rather than identifying a definitive endogenous induction mechanism, our results provide physiological evidence that experience-dependent population activity is accompanied by coordinated changes at both the network and synaptic levels in the hippocampus.

Acetylcholine (ACh) is a prominent neuromodulatory candidate that may facilitate experience-dependent high-frequency population activity in the hippocampus. Cholinergic signaling promotes synaptic plasticity in dorsal CA1, and both restraint stress and learning are accompanied by increased ACh release in this region in vivo (33,38–40). Moreover, the relationship between ACh release and contextual fear performance is abolished by muscarinic receptor blockade with scopolamine (41), and pharmacological inhibition of cholinergic signaling suppresses learning-associated synaptic plasticity in CA1 (33). Although our study does not directly manipulate cholinergic transmission, these findings are consistent with a modulatory role for ACh in enabling super burst activity and subsequent ripple firing diversification.

Although correlation alone cannot prove causation, the number and total duration of super burst events within the same animals were positively correlated with features of post-experience ripple firing, but not pre-experience ripple firing (Fig. 3H). In addition, emotionally salient experiences were accompanied by marked increases in super burst activity (Fig. 2), and these conditions were associated with strengthening of excitatory synaptic currents in CA1 pyramidal neurons (Fig. 4C). Together, these observations indicate a close relationship between experience-dependent population firing, subsequent ripple firing diversification, and synaptic reorganization.

Importantly, these relationships were observed selectively after experience, suggesting a temporal sequence in which super burst activity precedes changes in ripple firing features and synaptic inputs. While these data do not establish causality either, they are consistent with a model in which high-frequency population activity during emotionally salient experiences creates conditions that favor subsequent reorganization of ripple-associated firing patterns and synaptic efficacy. Such a sequence may provide a physiological substrate linking experience-dependent network dynamics with learning-related synaptic modifications.

In contrast to restraint stress and social interaction, exposure to a novel object selectively enhanced inhibitory synaptic currents, while excitatory synaptic parameters remained largely unchanged. Notably, however, increases in inhibitory synaptic strength and in no-firing silent periods were not restricted to the novel object condition. Across all four types of episodic experiences, both inhibitory postsynaptic currents (Fig. 4C) and the frequency of silent periods (Fig. 3B) showed a consistent post-experience increase.

One interpretation of this shared inhibitory enhancement is that diverse episodic experiences engage common network mechanisms that regulate overall excitability during memory-related processing. Although the behavioral and emotional content of the episodes differed, each experience contained elements of environmental novelty and behavioral salience, which may recruit inhibitory control in CA1 circuits. Consistent with this view, pharmacological manipulation of GABA_A_ receptors in CA1 has been shown to disrupt novel object memory when excessive inhibition is induced (42,43), while moderate suppression of background activity has been proposed to enhance signal discrimination and information throughput (44,45). Indeed, we recently demonstrated that inhibitory GABA_A_ synaptic plasticity is essential for contextual learning (46).

Inhibitory interneuron activity is known to be essential for both learning and memory processes and for the generation and temporal structuring of ripple oscillations (15,28,47). However, the causal relationship between strengthened inhibitory synaptic transmission and the emergence of silent periods remains unresolved. Experience-dependent enhancement of inhibitory synapses and silent periods may contribute to creating network conditions that support the precise timing and segregation of ripple-associated firing patterns.

SPW-Rs are also known to support replay of experience-dependent firing sequences, providing a physiological substrate for memory trace reactivation (48). While these studies establish the functional importance of SPW-Rs, most have focused on changes in ripple occurrence, duration, or replay content at the level of population averages. Although ripple firing reflects coordinated activity within CA1 local circuits, we did not directly monitor large-scale network dynamics in vivo. In contrast, our study demonstrates that individual ripple firing patterns exhibit experience-dependent diversification across multiple quantitative features, revealing fine-scale structure beyond population-level measures. By analyzing the fine structure of ripple-associated multi-unit firing, we show that recent episodic experience reshapes not only the frequency of ripple firing events but also the morphological and informational diversity of ripple firing, as reflected by increased feature dispersion and information entropy (Fig. 3E–G).

Importantly, this diversification was episode-type specific and persisted for at least 40 minutes after experience, indicating that ripple firing carries structured information about recent experience beyond simple replay. Given accumulating evidence that CA1 pyramidal neurons frequently exhibit dendritic plateau potentials and complex burst firing during learning (20), we specifically analyzed unsorted multi-unit activity in the 300–10000 Hz band to capture coordinated population firing underlying ripple generation. This approach allowed us to quantify experience-dependent reorganization of ripple firing patterns without spike sorting, which may overlook dynamic changes in spike waveform and synchrony during high-frequency events.

Our investigation of high-frequency activity surrounding episodic experiences revealed two distinct patterns of ripple activity modulation: experience-dependent global change and episode-specific reorganization across four quantitative features of ripple firing (amplitude, duration, arc length, and spike peaks; Figs. 3E-G). Comprehensive analysis of ripple firing similarity further revealed event-level specificity, indicating that ripple structure can encode recent experience (Fig. S4 and Table S9). This suggests that ripple firing may serve as a flexible population-level representational format, capable of incorporating information about recent experience through changes in their flexible structure.

The roles of excitatory and inhibitory synaptic transmission in SPW-R generation (49–51) and spike regulation during ripples are well established (15). In hippocampal slice models, SPW-Rs originate from CA3, propagate to CA1, and are critically dependent on AMPA receptor activation - their pharmacological blockade abolishes SPW-R generation (52,53). Inhibitory mechanisms involve two key processes: (i) CA3 interneurons coordinate pyramidal cell phase-locking through GABA_A_ receptor-mediated synchronous IPSCs (54), and (ii) pre-SWR inhibitory activity governs pyramidal cell spike timing and enables sequence pattern diversification (49). At the synaptic level, miniature events (mEPSCs/mIPSCs) reflect single vesicle glutamate/GABA release, with event frequency modulated by synaptic number or presynaptic release probability (55). Our study linked these findings to recent experience by demonstrating integrative plasticity at both synapse types, with two distinct patterns of synaptic modulation: experience-dependent change and episode-type specific reorganization (Fig. 4 and Table 3), providing synaptic evidence through which ripple firing diversity could be shaped by prior experience.

In support of this framework, preliminary observations from an independent cohort suggest that pharmacological blockade of muscarinic receptors with scopolamine prior to restraint stress suppresses the occurrence of super bursts, attenuates subsequent ripple firing diversification, and reduces freezing behavior in the same animals (56). Although these observations were not included in the present dataset and do not establish causality, they are consistent with the proposed role of cholinergic modulation in facilitating experience-dependent population activity and downstream plasticity.

Together, these findings position experience-dependent diversification of ripple firing—reflecting coordinated activity within CA1 local circuits—as a population-level signature coupled with corresponding reorganization of excitatory and inhibitory synaptic inputs onto CA1 pyramidal neurons. These coordinated changes suggest that episodic experience is not encoded at a single level of hippocampal organization, but rather through a structured sequence linking population activity, ripple firing reorganization, and synaptic plasticity.

### Our model consists of three stages

To address this gap, we propose a conceptual three-stage model (Fig. 5) that links experience-dependent population activity to persistent synaptic plasticity in hippocampal CA1 across multiple organizational levels. Rather than defining a complete mechanistic pathway, this model provides an organizing framework that integrates population activity, ripple firing reorganization, and synaptic plasticity observed in behaving animals.

**Figure 5.**
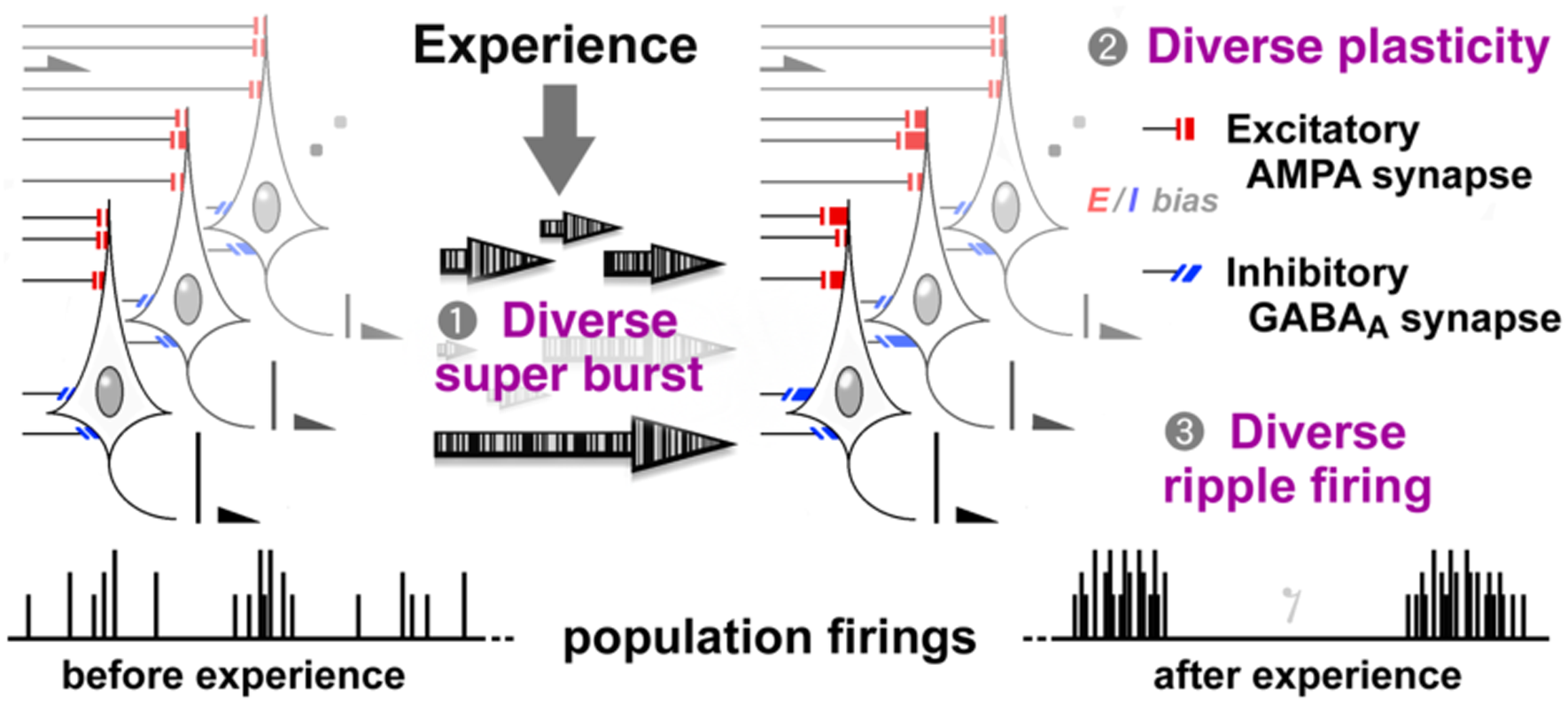
Conceptual three-stage model for experience-dependent encoding in hippocampal CA1. Distinct episodic experiences induce transient, experience-specific high-frequency population firing events “super bursts” in CA1 pyramidal neurons. These population-level events are proposed to bias experience-dependent diversification of excitatory AMPA and inhibitory GABA_A_ synaptic inputs, generating distinct synaptic configurations and biasing excitation–inhibition balance across CA1 microcircuits. In turn, these synaptic changes shape the diversification of ripple firing patterns, while strengthened inhibitory transmission contributes to the emergence of silent periods that temporally structure ripple activity. Through this coordinated cascade – from super burst activity to synaptic diversification and ripple firing reorganization – recent episodic experiences may be encoded within hippocampal CA1 circuits. Illustration is schematic and not drawn to scale; see Discussion for detailed rationale. The large number of CA1 pyramidal neurons (approximately 4 x 10^5^ in rats and 5 x 10^6^ in humans) underscores the population-level signature of this proposed encoding mechanism.

Our data indicate that experience-dependent super bursts exhibit early episode-type specificity in their temporal organization (Table 1). Based on this observation, we propose a cascading plasticity model in which these patterned population events bias subsequent ripple firing reorganization and establish experience-specific synaptic configurations through selective dendritic integration (Tables 2 and 3).

Although the causal relationships between experience, synaptic plasticity, and ripple dynamics remain to be fully established, several independent lines of evidence support the plausibility of this framework:

1. **Necessity of synaptic plasticity:** Single-unit in vivo recordings demonstrate that NMDA receptor–dependent plasticity is essential for encoding novel experience, as plasticity blockade impairs place field formation (57).
2. **Memory-specific reorganization:** Aversive stimuli induce rapid place cell remapping (≤ 5 min post-event), indicating experience-driven reconfiguration of network firing motifs (58).
3. **Experience-structured replay:** Large-scale ensemble analyses reveal that replay dynamics progress through structured stages (exploration → consolidation → retrieval) in an experience-dependent manner (59).

Consistent with this model, we recently demonstrated that inhibitory GABA_A_ synaptic plasticity is essential for contextual learning, that hippocampal commissural stimulation synchronized with super burst initiation blocks both learning and synaptic diversification, and that optogenetic inhibition of cholinergic input to the hippocampus selectively during contextual experience impairs learning in ChAT-Cre mice (46,60). These findings provide convergent causal support for a functional linkage between population activity, ripple reorganization, and integrative synaptic plasticity.

Together, these findings support an experience-dependent information processing pathway in which episode-specific population activity is transformed into a population-level signature through ripple firing reorganization, accompanied by coordinated synaptic plasticity at both excitatory and inhibitory synapses. As illustrated in Figure 5, this framework emphasizes temporal coordination across population activity, ripple firing reorganization, and synaptic plasticity as a key principle of hippocampal memory encoding. Future studies using temporally precise optogenetic or pharmacological interventions will be essential to directly test this model and explore its relevance to memory dysfunction.

## MATERIALS AND METHODS

### Animals

Male Sprague-Dawley rats (CLEA Japan, Tokyo, Japan) were housed individually at 24 ± 1°C under a 12-h light/dark cycle (lights on, 8:00–20:00) with ad libitum access to food (MF; Oriental Yeast Co., Tokyo, Japan). Animals aged 15–25 weeks were used for *in vivo* recordings. Episodic stimuli were provided by 8–15-week-old male or female rats, housed separately without electrodes. All procedures were approved by the Yamaguchi University Animal Care and Use Committee and were conducted in accordance with institutional guidelines and the NIH Guide for the Care and Use of Laboratory Animals.

### Surgery

Rats were anesthetized with sodium pentobarbital (50 mg/kg, i.p.) or a mixture of medetomidine (0.375 mg/kg), midazolam (2.0 mg/kg), and butorphanol (2.5 mg/kg). Then, they were positioned in a stereotaxic frame. Movable tungsten electrodes (50–80 kΩ; KS-216, Unique Medical Co., Japan) were chronically implanted just above the dorsal CA1 region of the hippocampus (AP: −3.0 to −3.6 mm; ML: ±1.4 to ±2.6 mm; DV: 2.0–2.2 mm) and secured with dental cement. Animals with histologically verified electrode placements outside CA1 were excluded from analysis.

### Multiple-unit Recording and Event Detection

Multiple-unit activity was recorded while animals freely behaved in their familiar home cages. Signals were amplified and transmitted via a headstage and shielded cable (MEG-2100 or MEG-6116; Nihon Kohden, Tokyo, Japan), band-pass filtered at 150–10,000 Hz, digitized at 25 kHz, and stored using Spike2 software (Cambridge Electronic Design, Cambridge, UK). All analyses focused on population spike activity (<u>300–10,000 Hz)</u>, and no conclusions were based on isolated single units unless explicitly stated.

### Definition of Super Burst

Super burst was defined as a population-level firing phenomenon characterized by a transient increase in multi-unit firing rate. Individual detected instances are referred to as super burst events. Events were identified when the firing rate exceeded +3 SD from baseline activity of low-noise period prior to experience. Super burst events typically did not exceed the threshold used for ripple detection (see below).

### Detection of Sharp-Wave Ripples and Ripple Firing

Only for the detection of sharp-wave ripples (SPW-Rs), the recorded signals were band-pass filtered at 150–300 Hz. The root mean square (RMS) of the filtered signal was calculated, and candidate SPW-R events were defined as epochs in which the RMS exceeded +6 SD above the baseline mean.

Ripple firing events were defined as short-duration, high-frequency multi-unit spike activity temporally associated with detected SPW-Rs and characterized by the presence of a peak complex. Individual ripple firing events had a mean duration of 56.3 ± 16.5 ms (mean ± SD; n = 5,333 events) and a signal-to-noise ratio ≥ 6:1. Signal segments corresponding to grooming or teeth grinding, which were characterized by symmetrical upper and lower peak amplitudes, were excluded from the analysis.

In contrast, super burst events did not meet the RMS threshold for SPW-R detection, lacked a peak complex, and exhibited longer event durations than ripple firing events (Fig. 1E). Behavioral states during ripple firing events were classified as eye-closed stationary (16.0 ± 7.0%), eye-open stationary (59.7 ± 11.0%), or eye-open moving (24.3 ± 11.3%; 1,631 events from 9 rats).

### Silent Periods

Silent periods were defined as inter-spike intervals exceeding +3 SD of the baseline distribution of inter-spike intervals.

### Spike Sorting and Visual Spike Classification

For illustrative purposes only (e.g., Fig. S1), spike sorting was performed using template-matching function in Spike2 software. Raw signals were high-pass filtered at 300 Hz, and candidate spikes were detected as threshold crossings with a signal-to-noise ratio ≥ 3:1. Spike waveform templates were generated using default Spike2 parameters. Template creation required at least eight spikes occurring at a frequency greater than 1 in 50 threshold-crossing events. Templates were generated with widths corresponding to 32% of the amplitude at each sampling point of the initial seed spike. At least 60% of the waveform was required to fall within the template, spanning from spike onset through the afterhyperpolarization period. Non-neuronal waveforms and artifacts were manually excluded by visual inspection.

Putative pyramidal neurons were defined by spike widths > 0.7 ms, and putative interneurons by widths < 0.4 ms (61). We performed these classifications solely to confirm that the recordings represented neuronal population activity.

### Behavioral Protocol and Recording Schedule

After ≥ 15 min of baseline recording in the home cage, animals were exposed for 10 min to one of four episodic conditions: restraint stress, social interaction with a female or male rat, or exposure to a novel object. Restraint stress involved gently tying the limbs with soft cloth and securing the animal to a wooden board (40). For the other experiences, a sexually mature female (postnatal age, 8–12 weeks), a young male (postnatal age, 6–7 weeks), or a novel object (yellow LEGO^®^/DUPLO^®^ brick; 15 × 8 × 3 cm) was placed in the home cage for 10 min. Neural recordings continued for ≥ 30 min following exposure. On the following day, animals were re-exposed to the same episode to assess experience-dependent effects (Fig. 1B).

### Histology

After completion of experiments, rats were deeply anesthetized with sodium pentobarbital (400 mg/kg, i.p.) and transcardially perfused with 0.1 M phosphate buffer containing 4% paraformaldehyde. Brains were post-fixed, cryoprotected in graded sucrose solutions (10–30%), and coronally sectioned at 40 µm. Sections were stained with hematoxylin and eosin, and electrode locations were verified using a rat brain atlas (62).

### Slice Patch-Clamp Recordings

Forty minutes after the onset of episodic exposure, animals were deeply anesthetized and coronal hippocampal slices (350 µm) containing CA1 were prepared using ice-cold dissection buffer and a vibratome (Leica VT-1200). Slices were incubated in oxygenated physiological solution (22–25 °C; 95% O₂/5% CO₂). Whole-cell voltage-clamp recordings were obtained from CA1 pyramidal neurons using patch pipettes (4–7 MΩ) filled with a cesium-based internal solution. Miniature excitatory and inhibitory postsynaptic currents (mEPSCs and mIPSCs) were sequentially recorded at −60 mV and 0 mV, respectively, in the presence of tetrodotoxin (0.5 µM). Event identity was confirmed pharmacologically using CNQX and bicuculline (10 µM each) (33,63,64,65).

### Entropy Analysis and Statistics

Appearance probabilities of ripple firing waveform features and synaptic parameters were estimated using one-dimensional kernel density estimation with bandwidth determined by Silverman’s rule (66,67). Shannon entropy was calculated in bits and log (1 + x)-transformed prior to parametric statistical analyses (68). Statistical comparisons employed one- and two-way ANOVA, repeated-measures ANOVA, MANOVA (Wilks’ lambda), and Fisher’s LSD post hoc tests. Spearman’s rank correlation was used to assess associations. Normality and homogeneity of variance were evaluated using Shapiro–Wilk and F tests, respectively. Statistical significance was set at *P* < 0.05.

## ACKNOWLEDGMENTS

The authors thank Drs. Sora Takayama, Koushi Seo, and Ryo Sato for the analysis of ripple firing events. This work was supported by Grant-in-Aid for Scientific Research B, 16H05129 (DM) and 19H03402 (DM), Grant-in-Aid for Scientific Research C, 26350988 (JI), 17K01987 (JI) and 25460314 (DM), and Scientific Research in Innovative Areas, 26115518 from the Ministry of Education, Culture, Sports, Science and Technology of Japan (DM). This project was also supported by YU AI project of center for information and data science education.

## AUTHOR CONTRIBUTIONS

DM and JI designed and performed the experiments. TT, JI, and DM analyzed firing events. DM and JI wrote the manuscript. DM organized the study, and all authors reviewed the manuscript.

## SUPPLEMENTAL INFORMATION

Supplemental information can be found online. It is on the server of Yamaguchi University Graduate School of Medicine.

- Movie S1
- Movie S2

## CONFLICT OF INTEREST

The authors declare no competing interests.

## Ethical statement

Our manuscript confirming the study is reported in accordance with ARRIVE guidelines (https://arriveguidelines.org).

## Data availability statement

Any additional information required to reanalyze the data reported in this paper is available from the lead contact upon request. Although the data are not publicly available due to the inclusion of data for ongoing research, all data will be uploaded to the web server of the Yamaguchi University Graduate School of Medicine upon acceptance of the research paper. In addition, all data will be deposited in an open repository.

## Notes

### Competing Interest Statement

The authors have declared no competing interest.

### Summary of Updates

We relocated Table 1 - 3 to the main body of this article. Those were mistakenly located in supplementary information.

