## Supplementary information for "Experience-dependent diversification of ripple firing and synaptic inputs in hippocampal CA1"

Figures S1 to S4,

Movies S1 and S2

**Table S1.** Temporal dynamics of silent periods across episodic experiences (related to Fig. 3B).

| Group | Overall | Time after onset | |
|---|---|---|---|
| | | Before *vs* 20 min | Before *vs* 40 min |
| Restraint | $F_{2, 16} = 3.740$, $P = 0.047$* | $P = 0.037$* | $P = 0.009$** |
| Female | $F_{2, 20} = 7.018$, $P = 0.005$** | $P = 0.131$ | $P = 0.006$** |
| Male | $F_{2, 20} = 8.557$, $P = 0.002$** | $P = 0.081$ | $P = 0.002$** |
| Object | $F_{2, 16} = 6.063$, $P = 0.011$* | $P = 0.054$ | $P = 0.018$* |
| Control | $F_{2, 12} = 0.534$, $P = 0.599$ | $P = 0.959$ | $P = 0.439$ |

ANOVA and post hoc analyses of changes in silent period duration across episodic experiences.

**Table S2.** Temporal dynamics of ripple firing across episodic experiences (related to Fig. 3C).

| Group | Overall | Time after onset | |
|---|---|---|---|
| | | Before *vs* 20 min | Before *vs* 40 min |
| Restraint | $F_{2, 16} = 18.672$, $P < 0.001$** | $P < 0.001$** | $P < 0.001$** |
| Female | $F_{2, 20} = 6.908$, $P = 0.005$** | $P = 0.023$* | $P = 0.004$** |
| Male | $F_{2, 20} = 4.992$, $P = 0.017$* | $P = 0.021$* | $P = 0.153$ |
| Object | $F_{2, 16} = 3.881$, $P = 0.042$* | $P = 0.196$ | $P = 0.052$ |
| Control | $F_{2, 12} = 1.991$, $P = 0.179$ | $P = 0.100$ | $P = 0.293$ |

ANOVA and post hoc analyses of changes in ripple firing incidence across episodic experiences.

**Table S3.** Temporal changes in ripple firing features following episodic experiences (related to Fig. 3E).

| Features | Experience | Time | Interaction |
|---|---|---|---|
| Amplitude | $F_{4, 5313} = 74.017$, $P < 0.001$** | $F_{3, 5313} = 31.614$, $P < 0.001$** | $F_{12, 5313} = 9.929$, $P < 0.001$** |
| Duration | $F_{4, 5313} = 47.086$, $P < 0.001$** | $F_{3, 5313} = 27.354$, $P < 0.001$** | $F_{12, 5313} = 7.638$, $P < 0.001$** |
| Arc length | $F_{4, 5313} = 86.803$, $P < 0.001$** | $F_{3, 5313} = 35.630$, $P < 0.001$** | $F_{12, 5313} = 12.565$, $P < 0.001$** |
| Peaks | $F_{4, 5313} = 154.271$, $P < 0.001$** | $F_{3, 5313} = 41.454$, $P < 0.001$** | $F_{12, 5313} = 21496$, $P < 0.001$** |

| Features | Episode | Overall | Time after onset | | |
| --- | --- | --- | --- | --- | --- |
| | | | 20 min | 30 min | 40 min |
| Amplitude | Restraint | $F_{3, 1039}$ = 21.104, $P$ < 0.001** | $P$ < 0.001** | $P$ < 0.001** | $P$ < 0.001** |
| | Female | $F_{3, 1255}$ = 6.314, $P$ < 0.001** | $P$ < 0.001** | $P$ < 0.001** | $P$ < 0.001** |
| | Male | $F_{3, 1226}$ = 16.251, $P$ < 0.001** | $P$ < 0.001** | $P$ < 0.001** | $P$ < 0.001** |
| | Object | $F_{3, 1016}$ = 1.991, $P$ = 0.114 | $P$ = 0.267 | $P$ = 0.841 | $P$ = 0.174 |
| | Control | $F_{3, 777}$ = 1.401, $P$ = 0.241 | $P$ = 0.284 | $P$ = 0.041 | $P$ = 0.298 |
| Duration | Restraint | $F_{3, 1039}$ = 23.366, $P$ < 0.001** | $P$ < 0.001** | $P$ < 0.001** | $P$ < 0.001** |
| | Female | $F_{3, 1255}$ = 14.112, $P$ < 0.001** | $P$ = 0.030* | $P$ < 0.001** | $P$ < 0.001** |
| | Male | $F_{3, 1226}$ = 11.690, $P$ < 0.001** | $P$ < 0.001** | $P$ < 0.001** | $P$ < 0.001** |
| | Object | $F_{3, 1016}$ = 2.770, $P$ = 0.041* | $P$ = 0.051 | $P$ = 0.395 | $P$ = 0.969 |
| | Control | $F_{3, 777}$ = 0.283, $P$ = 0.838 | $P$ = 0.969 | $P$ = 0.678 | $P$ = 0.450 |
| Arc length | Restraint | $F_{3, 1039}$ = 30.629, $P$ < 0.001** | $P$ < 0.001** | $P$ < 0.001** | $P$ < 0.001** |
| | Female | $F_{3, 1255}$ = 15.187, $P$ < 0.001** | $P$ < 0.001** | $P$ < 0.001** | $P$ < 0.001** |
| | Male | $F_{3, 1226}$ = 14.681, $P$ < 0.001** | $P$ < 0.001** | $P$ < 0.001** | $P$ < 0.001** |
| | Object | $F_{3, 1016}$ = 3.616, $P$ = 0.013* | $P$ = 0.075 | $P$ = 0.129 | $P$ = 0.780 |
| | Control | $F_{3, 777}$ = 0.212, $P$ = 0.888 | $P$ = 0.945 | $P$ = 0.884 | $P$ = 0.515 |
| Peaks | Restraint | $F_{3, 1039}$ = 34.404, $P$ < 0.001** | $P$ < 0.001** | $P$ < 0.001** | $P$ < 0.001** |
| | Female | $F_{3, 1255}$ = 19.804, $P$ < 0.001** | $P$ < 0.001** | $P$ < 0.001** | $P$ < 0.001** |
| | Male | $F_{3, 1226}$ = 32.537, $P$ < 0.001** | $P$ < 0.001** | $P$ < 0.001** | $P$ < 0.001** |
| | Object | $F_{3, 1016}$ = 17.392, P < 0.001** | $P$ = 0.009** | $P$ < 0.001** | $P$ < 0.001** |
| | Control | $F_{3, 777}$ = 0.083, P = 0.969 | $P$ = 0.816 | $P$ = 0.827 | $P$ =0.862 |

Two-way repeated-measures ANOVA and post hoc analyses of changes in individual ripple firing features (amplitude, duration, arc length, and number of negative peaks) across episodic experiences.

**Table S4.** Temporal changes in ripple firing self-entropies following episodic experiences (related to Fig. 3F).

| Features | Experience | Time | Interaction |
| --- | --- | --- | --- |
| Amplitude | $F_{4, 5313}$ = 84.791, $P$ < 0.001** | $F_{3, 5313}$ = 28.665, $P$ < 0.001** | $F_{12, 5313}$ = 9.744, $P$ < 0.001** |

| | | | |
|---|---|---|---|
| Duration | $F_{4, 5313} = 39.091$, $P < 0.001$** | $F_{3, 5313} = 24.131$, $P < 0.001$** | $F_{12, 4536} = 3.616$, $P < 0.001$** |
| Arc length | $F_{4, 5313} = 72.606$, $P < 0.001$** | $F_{3, 5313} = 26.149$, $P < 0.001$** | $F_{12, 4536} = 7.867$, $P < 0.001$** |
| Peaks | $F_{4, 5313} = 94.283$, $P < 0.001$** | $F_{3, 5313} = 26.992$, $P < 0.001$** | $F_{12, 4536} = 10.312$, $P < 0.001$** |

| | | | Time after onset | | |
|---|---|---|---|---|---|
| Features | Episode | Overall | 20 min | 30 min | 40 min |
| Amplitude | Restraint | $F_{3, 1039} = 14.789$, $P < 0.001$** | $P < 0.001$** | $P < 0.001$** | $P < 0.001$** |
| | Female | $F_{3, 1255} = 7.513$, $P < 0.001$** | $P < 0.001$** | $P < 0.001$** | $P < 0.001$** |
| | Male | $F_{3, 1226} = 14.822$, $P < 0.001$** | $P = 0.002$** | $P < 0.001$** | $P < 0.001$** |
| | Object | $F_{3, 1016} = 8.336$, $P < 0.001$** | $P < 0.001$** | $P = 0.013$* | $P < 0.001$** |
| | Control | $F_{3, 777} = 0.437$, $P = 0.726$ | $P = 0.258$ | $P = 0.484$ | $P = 0.574$ |
| Duration | Restraint | $F_{3, 1039} = 13.618$, $P < 0.001$** | $P < 0.001$** | $P < 0.001$** | $P < 0.001$** |
| | Female | $F_{3, 1255} = 3.351$, $P = 0.018$* | $P = 0.065$ | $P = 0.114$ | $P = 0.002$** |
| | Male | $F_{3, 1226} = 4.174$, $P = 0.006$** | $P = 0.002$** | $P = 0.030$* | $P = 0.003$** |
| | Object | $F_{3, 1016} = 4.218$, $P = 0.006$** | $P < 0.001$** | $P = 0.229$ | $P = 0.065$ |
| | Control | $F_{3, 777} = 2.581$, $P = 0.052$ | $P = 0.075$ | $P = 0.113$ | $P = 0.007$ |
| Arc length | Restraint | $F_{3, 1039} = 29.320$, $P < 0.001$** | $P < 0.001$** | $P < 0.001$** | $P < 0.001$** |
| | Female | $F_{3, 1255} = 3.889$, $P = 0.009$** | $P = 0.034$* | $P = 0.294$ | $P < 0.001$** |
| | Male | $F_{3, 1226} = 4.780$, $P = 0.003$** | $P < 0.001$** | $P = 0.008$** | $P < 0.001$** |
| | Object | $F_{3, 1016} = 2.368$, $P = 0.069$ | $P = 0.009$** | $P = 0.297$ | $P = 0.009$** |
| | Control | $F_{3, 777} = 0.198$, $P = 0.898$ | $P = 0.450$ | $P = 0.806$ | $P = 0.762$ |
| Peaks | Restraint | $F_{3, 1039} = 16.020$, $P < 0.001$** | $P < 0.001$** | $P < 0.001$** | $P < 0.001$** |
| | Female | $F_{3, 1255} = 6.188$, $P < 0.001$** | $P = 0.039$* | $P < 0.001$** | $P < 0.001$** |
| | Male | $F_{3, 1226} = 11.113$, $P < 0.001$** | $P < 0.001$** | $P < 0.001$** | $P < 0.001$** |
| | Object | $F_{3, 1016} = 3.404$, $P = 0.017$* | $P = 0.103$ | $P = 0.113$ | $P < 0.001$** |
| | Control | $F_{3, 777} = 0.125$, $P = 0.945$ | $P = 0.842$ | $P = 0.741$ | $P = 0.837$ |

Two-way repeated-measures ANOVA and post hoc analyses of changes in self-entropy of individual ripple firing features across episodic experiences.

**Table S5.** Detailed MANOVA results for integrated ripple firing features (related to Fig. 3G).

| Comparison | Experience | Time | Interaction |
|---|---|---|---|
| Overall | $F_{12,\ 11993}$ = 48.996 $P$ < 0.001** | $F_{12,\ 11993}$ = 21.455 $P$ < 0.001** | $F_{36,\ 16989}$ = 10.206 P < 0.001** |
| Restraint *vs* Female | $F_{4,\ 2291}$ = 30.810 $P$ < 0.001** | $F_{12,\ 6062}$ = 21.834 $P$ < 0.001** | $F_{12,\ 6062}$ = 7.905 $P$ < 0.001** |
| Restraint *vs* Male | $F_{4,\ 2262}$ = 31.515 $P$ < 0.001** | $F_{12,\ 5985}$ = 23.057 $P$ < 0.001** | $F_{12,\ 5985}$ = 8.125 $P$ < 0.001** |
| Restraint *vs* Object | $F_{4,\ 2052}$ = 86.412 $P$ < 0.001** | $F_{12,\ 5429}$ = 10.448 $P$ < 0.001** | $F_{12,\ 5429}$ = 16.409 $P$ < 0.001** |
| Restraint *vs* Control | $F_{4,\ 1813}$ = 48.846 $P$ < 0.001** | $F_{12,\ 4797}$ = 9.791 $P$ < 0.001** | $F_{12,\ 4797}$ = 9.808 $P$ < 0.001** |
| Female *vs* Male | $F_{4,\ 2478}$ = 3.395 $P$ = 0.009** | $F_{12,\ 6556}$ = 17.446 $P$ < 0.001** | $F_{12,\ 6556}$ = 1.016 $P$ = 0.431 |
| Female *vs* Object | $F_{4,\ 2268}$ = 60.959 $P$ < 0.001** | $F_{12,\ 6001}$ = 3.988 $P$ < 0.001** | $F_{12,\ 6001}$ = 10.322 $P$ < 0.001** |
| Female *vs* Control | $F_{4,\ 2029}$ = 18.258 $P$ < 0.001** | $F_{12,\ 5369}$ = 3.962 $P$ < 0.001** | $F_{12,\ 5369}$ = 3.847 $P$ < 0.001** |
| Male *vs* Object | $F_{4,\ 2239}$ = 94.403 $P$ < 0.001** | $F_{12,\ 5924}$ = 4.304 $P$ < 0.001** | $F_{12,\ 5924}$ = 13.257 $P$ < 0.001** |
| Male *vs* Control | $F_{4,\ 2000}$ = 32.546 $P$ < 0.001** | $F_{12,\ 5292}$ = 5.224 $P$ < 0.001** | $F_{12,\ 5292}$ = 4.849 $P$ < 0.001** |
| Object *vs* Control | $F_{4,\ 1790}$ = 23.418 $P$ < 0.001** | $F_{12,\ 4736}$ = 4.662 $P$ < 0.001** | $F_{12,\ 4736}$ = 4.405, $P$ < 0.001** |

| Comparison | 20 min | 30 min | 40 min |
|---|---|---|---|
| Overall | $F_{12,\ 2948}$ = 13.833 $P$ < 0.001** | $F_{12,\ 2953}$ = 25.382, $P$ < 0.001** | $F_{12,\ 2974}$ = 20.724, $P$ < 0.001** |
| Restraint *vs* Female | $F_{4,\ 570}$ = 8.723, $P$ < 0.001** | $F_{4,\ 553}$ = 19.624, $P$ < 0.001** | $F_{4,\ 579}$ = 9.331, $P$ < 0.001** |
| Restraint *vs* Male | $F_{4,\ 545}$ = 11.160, $P$ < 0.001** | $F_{4,\ 555}$ = 21.629, $P$ < 0.001** | $F_{4,\ 563}$ = 9.292, $P$ < 0.001** |
| Restraint *vs* Object | $F_{4,\ 493}$ = 21.132, $P$ < 0.001** | $F_{4,\ 518}$ = 39.137, $P$ < 0.001** | $F_{4,\ 500}$ = 36.066, $P$ < 0.001** |
| Restraint *vs* Control | $F_{4,\ 441}$ = 14.056 $P$ < 0.001** | $F_{4,\ 443}$ = 21.966 $P$ < 0.001** | $F_{4,\ 458}$ = 14.193 $P$ < 0.001** |
| Female *vs* Male | $F_{4,\ 618}$ = 2.642, $P$ = 0.033* | $F_{4,\ 595}$ = 1.367, $P$ = 0.244 | $F_{4,\ 621}$ = 1.465, $P$ = 0.211 |

| | | | |
|---|---|---|---|
| Female *vs* Object | $F_{4, 566} = 14.587$, $P < 0.001$** | $F_{4, 558} = 24.556$, $P < 0.001$** | $F_{4, 567} = 14.627$, $P < 0.001$** |
| Female *vs* Control | $F_{4, 514} = 3.782$ $P = 0.005$** | $F_{4, 483} = 8.296$ $P < 0.001$** | $F_{4, 516} = 15.084$ $P < 0.001$** |
| Male *vs* Object | $F_{4, 541} = 23.424$, $P < 0.001$** | $F_{4, 560} = 41.345$, $P < 0.001$** | $F_{4, 542} = 57.100$, $P < 0.001$** |
| Male *vs* Control | $F_{4, 489} = 8.247$ $P < 0.001$** | $F_{4, 485} = 17.859$ $P < 0.001$** | $F_{4, 500} = 16.781$ $P < 0.001$** |
| Object *vs* Control | $F_{4, 437} = 8.771$ $P < 0.001$** | $F_{4, 448} = 6.983$ $P < 0.001$** | $F_{4, 437} = 19.668$ $P < 0.001$** |

| Episode | Overall | before *vs* 20 min | before *vs* 30 min | before *vs* 40 min |
|---|---|---|---|---|
| Restraint | $F_{12, 2741} = 14.315$ $P < 0.001$** | $F_{4, 516} = 17.976$ $P < 0.001$** | $F_{4, 524} = 29.407$ $P < 0.001$** | $F_{4, 528} = 24.036$ $P < 0.001$** |
| Female | $F_{12, 3313} = 8.058$ $P < 0.001$** | $F_{4, 634} = 5.772$ $P < 0.001$** | $F_{4, 609} = 12.089$ $P < 0.001$** | $F_{4, 631} = 19.797$ $P < 0.001$** |
| Male | $F_{12, 3236} = 10.286$ $P < 0.001$** | $F_{4, 619} = 12.169$ $P < 0.001$** | $F_{4, 621} = 24.168$ $P < 0.001$** | $F_{4, 625} = 24.736$ $P < 0.001$** |
| Object | $F_{12, 2680} = 7.477$ $P < 0.001$** | $F_{4, 509} = 8.469$ $P < 0.001$** | $F_{4, 526} = 8.056$ $P < 0.001$** | $F_{4, 504} = 17.715$ $P < 0.001$** |
| Control | $F_{12, 2048} = 0.843$ $P = 0.606$ | $F_{4, 387} = 0.502$ $P = 0.734$ | $F_{4, 381} = 1.947$ $P = 0.102$ | $F_{4, 392} = 0.763$ $P = 0.550$ |

Detailed MANOVA and post hoc results for integrated ripple firing features (amplitude, duration, arc length, and number of negative peaks) summarized in Table 2.

**Table S6.** Detailed MANOVA results for integrated ripple firing self-entropies.

| Comparison | Experience | Time | Interaction |
|---|---|---|---|
| Overall | $F_{16, 16223}$ = 35.748<br>$P$ < 0.001** | $F_{12, 14049}$ = 13.595<br>$P$ < 0.001** | $F_{48, 20457}$ = 4.565<br>$P$ < 0.001** |
| Restraint *vs* Female | $F_{4, 2291}$ = 33.374<br>$P$ < 0.001** | $F_{12, 6062}$ = 11.391<br>$P$ < 0.001** | $F_{12, 6062}$ = 6.377<br>$P$ < 0.001** |
| Restraint *vs* Male | $F_{4, 2262}$ = 42.701<br>$P$ < 0.001** | $F_{12, 5985}$ = 12.840<br>$P$ < 0.001** | $F_{12, 5985}$ = 5.364<br>$P$ < 0.001** |
| Restraint *vs* Object | $F_{4, 2052}$ = 48.154<br>$P$ < 0.001** | $F_{12, 5429}$ = 9.476<br>$P$ < 0.001** | $F_{12, 5429}$ = 6.414<br>$P$ < 0.001** |
| Restraint *vs* Control | $F_{4, 1813}$ = 52.735<br>$P$ < 0.001** | $F_{12, 4797}$ = 6.922<br>$P$ < 0.001** | $F_{12, 4797}$ = 5.335<br>$P$ < 0.001** |
| Female *vs* Male | $F_{4, 2478}$ = 3.920<br>$P$ = 0.004** | $F_{12, 6556}$ = 10.656<br>$P$ < 0.001** | $F_{12, 6556}$ = 0.736<br>$P$ = 0.717 |
| Female *vs* Object | $F_{4, 2268}$ = 12.403<br>$P$ < 0.001** | $F_{12, 6001}$ = 6.955<br>$P$ = 0.010* | $F_{12, 6001}$ = 1.530<br>$P$ = 0.106 |
| Female *vs* Control | $F_{4, 2029}$ = 43.749<br>$P$ < 0.001** | $F_{12, 5369}$ = 3.851<br>$P$ < 0.001** | $F_{12, 5369}$= 2.186<br>$P$ = 0.010* |
| Male *vs* Object | $F_{4, 2239}$ = 6.812<br>$P$ < 0.001** | $F_{12, 5924}$ = 8.355<br>$P$ < 0.001** | $F_{12, 5924}$ = 2.023<br>$P$ = 0.019* |
| Male *vs* Control | $F_{4, 2000}$ = 31.836<br>$P$ < 0.001** | $F_{12, 5252}$ = 4.669<br>$P$ < 0.001** | $F_{12, 5292}$= 3.054<br>$P$ < 0.001** |
| Object *vs* Control | $F_{4, 1790}$ = 29.895<br>$P$ < 0.001** | $F_{12, 4736}$ = 3.432<br>$P$ < 0.001** | $F_{12, 4736}$= 1.500<br>$P$ = 0.116 |

| Comparison | 20 min | 30 min | 40 min |
|---|---|---|---|
| Overall | $F_{16, 3997}$ = 8.602<br>$P$ < 0.001** | $F_{12, 3984}$ = 15.047<br>$P$ < 0.001** | $F_{16, 4042}$ = 11.964<br>$P$ < 0.001** |
| Restraint *vs* Female | $F_{4, 570}$ = 7.994<br>$P$ < 0.001** | $F_{4, 553}$ = 16.972<br>$P$ < 0.001** | $F_{4, 579}$ = 11.892<br>$P$ < 0.001** |
| Restraint *vs* Male | $F_{4, 545}$ = 9.404<br>$P$ < 0.001** | $F_{4, 555}$ = 18.128<br>$P$ < 0.001** | $F_{4, 563}$ = 14.845<br>$P$ < 0.001** |
| Restraint *vs* Object | $F_{4, 493}$ = 10.872<br>$P$ < 0.001** | $F_{4, 518}$ = 22.529<br>$P$ < 0.001** | $F_{4, 500}$ = 17.241<br>$P$ < 0.001** |
| Restraint *vs* Control | $F_{4, 441}$ = 12.262<br>$P$ < 0.001** | $F_{4, 443}$ = 20.815<br>$P$ < 0.001** | $F_{4, 458}$ = 20.400<br>$P$ < 0.001** |

| | | | |
|---|---|---|---|
| Female *vs* Male | $F_{4, 618} = 1.553$ $P = 0.185$ | $F_{4, 595} = 0.958$ $P = 0.430$ | $F_{4, 621} = 1.460$ $P = 0.213$ |
| Female *vs* Object | $F_{4, 566} = 3.273$ $P = 0.011$* | $F_{4, 558} = 5.254$ $P < 0.001$** | $F_{4, 558} = 5.151$ $P < 0.001$** |
| Female *vs* Control | $F_{4, 514} = 8.520$ $P < 0.001$** | $F_{4, 483} = 17.959$ $P < 0.001$** | $F_{4, 516} = 12.361$ $P < 0.001$** |
| Male *vs* Object | $F_{4, 541} = 4.238$ $P = 0.002$** | $F_{4, 560} = 4.748$ $P < 0.001$** | $F_{4, 542} = 1.033$ $P = 0.390$ |
| Male *vs* Control | $F_{4, 489} = 7.348$ $P < 0.001$** | $F_{4, 485} = 15.743$ $P < 0.001$** | $F_{4, 500} = 7.867$ $P < 0.001$** |
| Object *vs* Control | $F_{4, 437} = 8.491$ $P < 0.001$** | $F_{4, 448} = 8.148$ $P < 0.001$** | $F_{4, 437} = 9.831$ $P < 0.001$** |

| Comparison | Overall | before *vs* 20 min | before *vs* 30 min | before *vs* 40 min |
|---|---|---|---|---|
| Restraint | $F_{12, 2741} = 9.300$, $P < 0.001$** | $F_{4, 516} = 14.037$, $P < 0.001$** | $F_{4, 524} = 24.082$, $P < 0.001$** | $F_{4, 528} = 24.446$, $P < 0.001$** |
| Female | $F_{12, 3313} = 5.084$, $P < 0.001$** | $F_{4, 634} = 5.597$, $P < 0.001$** | $F_{4, 609} = 13.379$, $P < 0.001$** | $F_{4, 631} = 11.518$, $P < 0.001$** |
| Male | $F_{12, 3236} = 6.247$, $P < 0.001$** | $F_{4, 619} = 10.190$, $P < 0.001$** | $F_{4, 621} = 17.219$, $P < 0.001$** | $F_{4, 625} = 14.331$, $P < 0.001$** |
| Object | $F_{12, 2680} = 3.843$, $P < 0.001$** | $F_{4, 509} = 9.221$, $P < 0.001$** | $F_{4, 526} = 3.108$, $P = 0.015$* | $F_{4, 504} = 7.621$, $P < 0.001$** |
| Control | $F_{12, 2048} = 1.370$ $P = 0.173$ | $F_{4, 387} = 2.772$ $P = 0.027$ | $F_{4, 381} = 2.997$ $P = 0.019$ | $F_{4, 392} = 2.984$ $P = 0.019$ |

Detailed MANOVA and post hoc results for integrated self-entropy of ripple firing summarized in Table 2.

**Table S7.** Synaptic input properties across episodic experiences (related to Fig. 4C).

| Comparison | mEPSC amplitude (pA) | mIPSC amplitude (pA) | mEPSC frequency (5 $min^{-1}$) | mIPSC frequency (5 $min^{-1}$) |
|---|---|---|---|---|
| Overall | $F_{4,183} = 9.442$, $P < 0.001$** | $F_{4, 183} = 5.227$, $P < 0.001$** | $F_{4, 183} = 2.869$, $P = 0.025$* | $F_{4, 183} = 7.955$, $P < 0.001$** |
| Control *vs* Restraint | $P < 0.001$** | $P = 0.043$* | $P = 0.069$ | $P = 0.274$ |
| Control *vs* Female | $P < 0.001$** | $P = 0.049$* | $P = 0.022$* | $P = 0.703$ |
| Control *vs* Male | $P < 0.001$** | $P < 0.001$** | $P = 0.024$* | $P < 0.001$** |
| Control *vs* Object | $P = 0.370$ | $P = 0.012$* | $P = 0.887$ | $P = 0.464$ |

ANOVA of excitatory and inhibitory synaptic input parameters recorded ex vivo from CA1 pyramidal neurons following episodic experiences.

**Table S8.** Entropy analysis of synaptic input properties (related to Fig. 4E).

| Comparison | mEPSC amplitude (bit) | mIPSC amplitude (bit) | mEPSC frequency (bit) | mIPSC frequency (bit) |
|---|---|---|---|---|
| Overall | $F_{4,183} = 4.057$, $P = 0.004$** | $F_{4, 183} = 4.381$, $P = 0.002$** | $F_{4, 183} = 2.723$, $P = 0.031$* | $F_{4, 183} = 8.431$, $P < 0.001$** |
| Control *vs* Restraint | $P = 0.023$* | $P = 0.124$ | $P = 0.447$ | $P = 0.296$ |
| Control *vs* Female | $P < 0.001$** | $P = 0.013$* | $P = 0.226$ | $P = 0.739$ |
| Control *vs* Male | $P = 0.023$* | $P < 0.001$** | $P = 0.018$* | $P < 0.001$** |
| Control *vs* Object | $P = 0.692$ | $P = 0.105$ | $P = 0.459$ | $P = 0.026$* |

ANOVA of information entropy of excitatory and inhibitory synaptic inputs parameters following episodic experiences.

**Table S9.** Temporal changes in the Euclidean distance between ripple firing waveforms following episodic experiences (related to Fig. S4).

| Comparison | Experience | Time | Interaction |
|---|---|---|---|
| Overall | $F_{4, 82581}$ = 995.96<br>$P$ < 0.001** | $F_{3, 82581}$ = 548.44<br>$P$ < 0.001** | $F_{12, 82581}$ = 138.66<br>$P$ < 0.001** |
| Restraint *vs* Female | $F_{1, 36544}$ = 1170.6<br>$P$ < 0.001** | $F_{3, 36544}$ = 439.48<br>$P$ < 0.001** | $F_{3, 36544}$ = 158.51<br>$P$ < 0.001** |
| Restraint *vs* Male | $F_{1, 35282}$ = 1670.7<br>$P$ < 0.001** | $F_{3, 35282}$ = 494.34<br>$P$ < 0.001** | $F_{3, 35282}$ = 218.27<br>$P$ < 0.001** |
| Restraint *vs* Object | $F_{1, 32378}$ = 1427.4<br>$P$ < 0.001** | $F_{3, 32378}$ = 341.58<br>$P$ < 0.001** | $F_{3, 32378}$ = 200.81<br>$P$ < 0.001** |
| Restraint *vs* Control | $F_{1, 27019}$ = 1205.3<br>$P$ < 0.001** | $F_{3, 27019}$ = 248.37<br>$P$ < 0.001** | $F_{3, 27019}$ = 152.15<br>$P$ < 0.001** |
| Female *vs* Male | $F_{1, 39338}$ = 44.113<br>$P$ < 0.001** | $F_{3, 39338}$ = 233.38<br>$P$ < 0.001** | $F_{3, 39338}$ = 28.027<br>$P$ < 0.001** |
| Female *vs* Object | $F_{1, 36434}$ = 70.428<br>$P$ < 0.001** | $F_{3, 36434}$ = 124.27<br>$P$ < 0.001** | $F_{3, 36434}$ = 12.074<br>$P$ < 0.001** |
| Female *vs* Control | $F_{1, 31165}$ = 124.08<br>$P$ < 0.001** | $F_{3, 31165}$ = 94.821<br>$P$ < 0.001** | $F_{3, 31165}$ = 14.246<br>$P$ < 0.001** |
| Male *vs* Object | $F_{1, 35172}$ = 16.890<br>$P$ < 0.001** | $F_{3, 35172}$ = 184.90<br>$P$ < 0.001** | $F_{3, 36172}$ = 71.489<br>$P$ < 0.001** |
| Male *vs* Control | $F_{1, 29903}$ = 127.86<br>$P$ < 0.001** | $F_{3, 29903}$ = 317.53<br>$P$ < 0.001** | $F_{3, 29903}$ = 60.647<br>$P$ < 0.001** |
| Object *vs* Control | $F_{1, 26999}$ = 14.564<br>$P$ < 0.001** | $F_{3, 26999}$ = 79.649<br>$P$ < 0.001** | $F_{3, 26999}$ = 11.292<br>$P$ < 0.001** |

Two-way repeated-measures ANOVA and post hoc analyses of changes in Euclidean distances from 82,601 pairs of ripple firing waveforms.

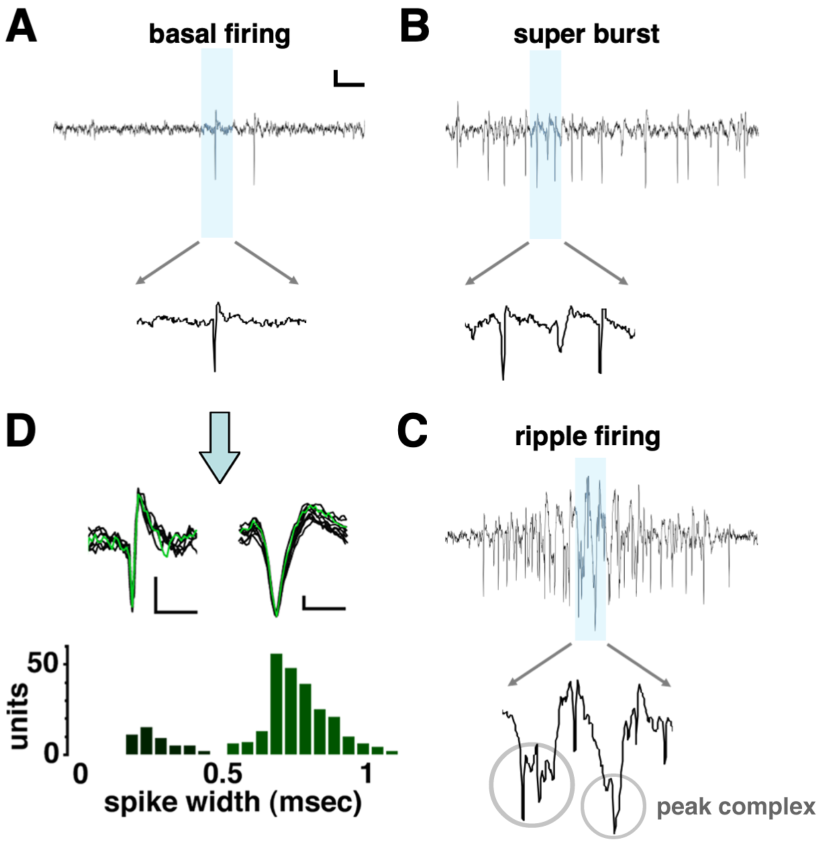


**Figure S1. Limitations of spike classification during super bursts and ripple firing events. (A–C)** Representative examples of CA1 multi-unit activity recorded at 25 kHz and filtered between 300 - 10000 Hz. Shaded regions indicate the time window used for waveform extraction. (**A**) Basal firing, showing isolated spike waveforms suitable for conventional spike width-based classification. Scale bar = 10 ms. (**B**) Super burst, in which overlapping and distorted spike waveforms prevent reliable template matching. (**C**) Ripple firing event, characterized by high-frequency oscillatory activity accompanied by complex multi-peak waveforms ("peak complexes"), precluding identification of single-unit spikes. (**D**) Spike width distribution obtained from basal firing periods. Overlaid waveforms illustrate narrow- and broad-spike populations, corresponding to putative interneurons and pyramidal neurons, respectively. Histogram shows that spike width-based classification is feasible during basal firing but becomes unreliable during super bursts and ripple firing events due to waveform distortion and temporal overlap. Scale bars: vertical, 0.2 μV; horizontal, 1 ms.

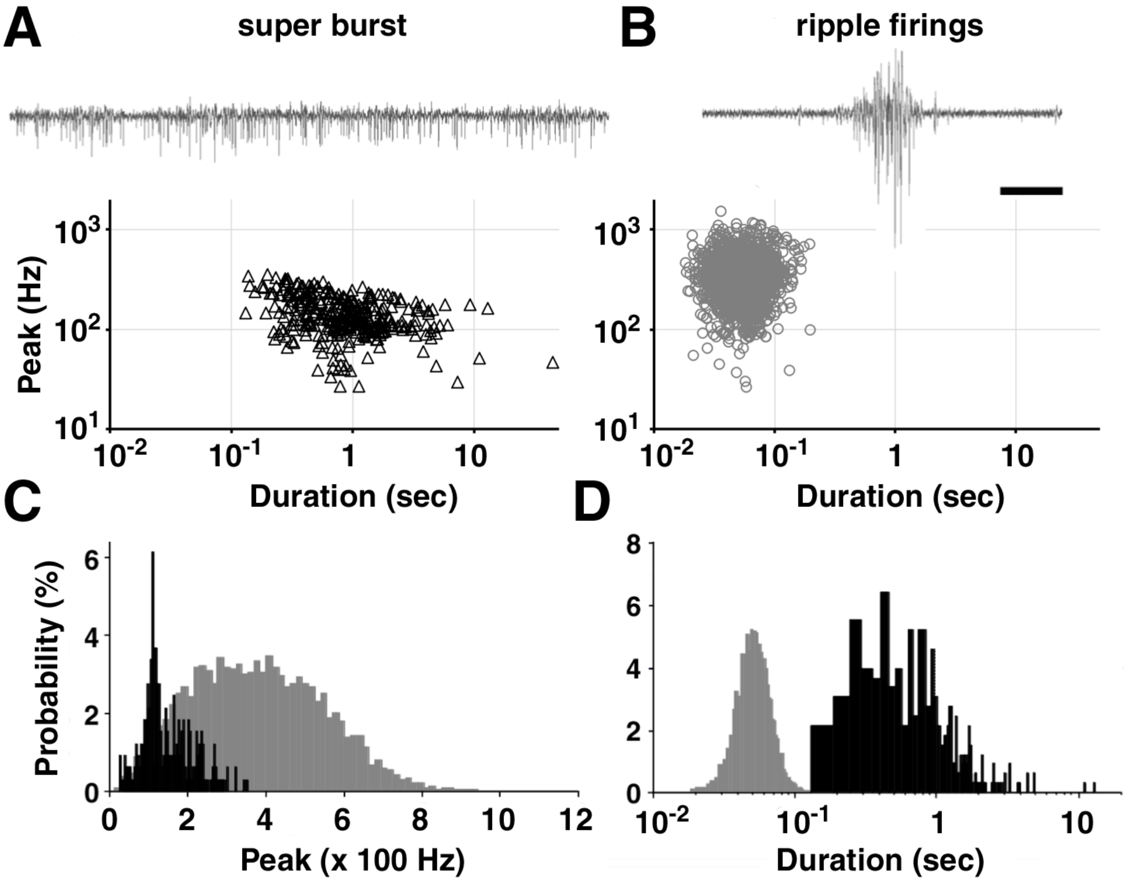


**Figure S2. Two extracted features of super bursts and ripple firing. (A)** An example trace of a super burst event. The peak frequency (Hz) and duration of the super burst were plotted for individual events (N = 327). **(B)** An example trace of a ripple firing event. The peak frequency (Hz) and duration of the event were plotted across events (N = 5333). Scale bar = 50 ms. **(C)** Histogram of peak frequency for ripple firing (gray) and super bursts (black). **(D)** Histogram of duration for ripple firing (gray) and super bursts (black).

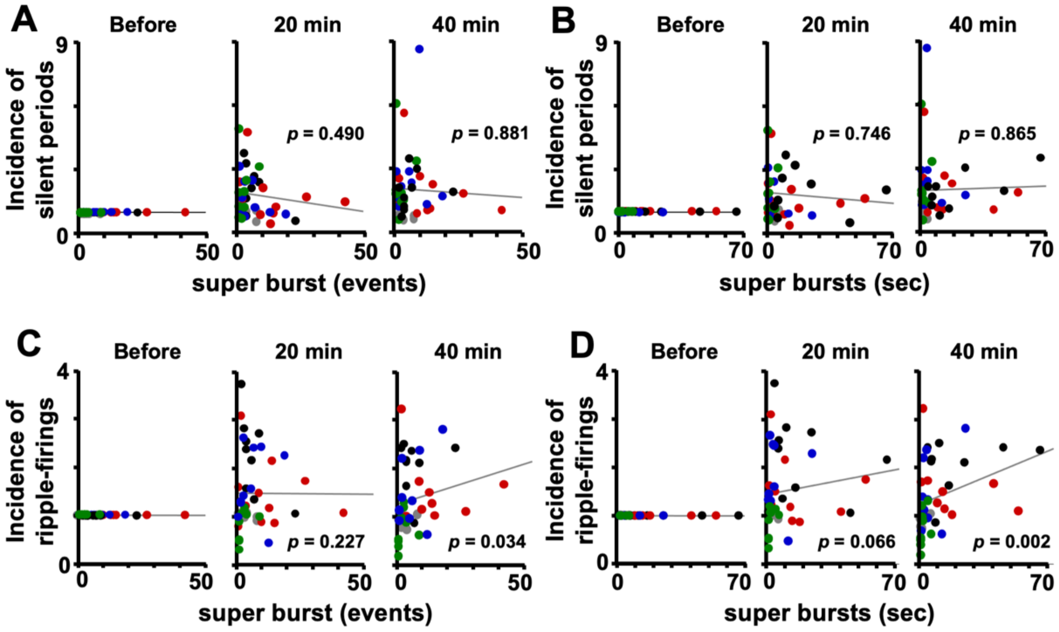


**Figure S3. Super bursts correlate with ripple firing, but not with silent periods.**
**(A–B)** The increase in silent periods was not significantly correlated with the number (**A**) or total duration (**B**) of super bursts. **(C–D)** In contrast, the number (**C**) and total duration (**D**) of super bursts were significantly correlated with the increase in ripple firing incidence. Each plot shows data from individual animals, normalized to their pre-experience baseline. Colors indicate episode type: restraint stress (black), contact with female (red), contact with male (blue), novel object (green), control (gray).

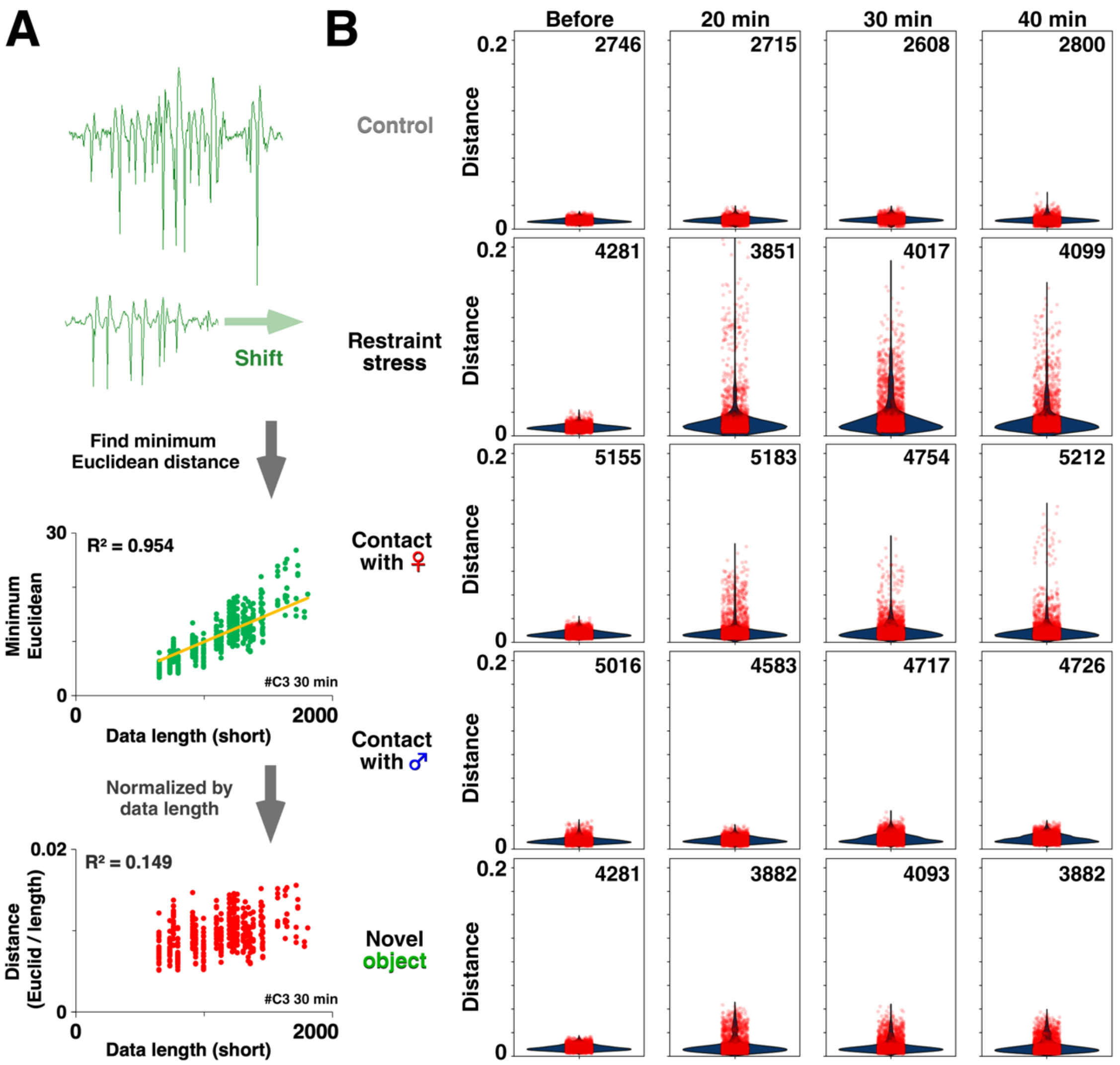


**Figure S4. Ripple firing waveforms exhibit episode-specific differences in event-level similarity.** (A) Method for comparing individual ripple firing waveforms. For each pair of ripple firing events, waveforms were temporally shifted to minimize Euclidean distance and normalized to account for differences in event duration. (B) Violin plots showing the distribution of minimum Euclidean distances between all possible pairs of ripple firing events before and after episodic experience (red). Each panel indicates the total number of ripple firing pairs analyzed, together with kernel density estimates (blue). Identical ripple firing waveforms would yield a distance of zero; however, no two ripple firing events were identical. Importantly, this single-parameter metric — the Euclidean distance between event-level ripple firing waveforms — captured both episode-dependent shifts and episode-specific convergence in waveform similarity following experience (see Table S9 for statistics).

**Movie S1. (separate file) Pre-experience CA1 multi-unit activity in a freely moving rat.** This movie displays multi-unit neuronal activity recorded from the hippocampal CA1 region before the onset of episodic experience (150–10,000 Hz). The background audio corresponds to spike firing. During the baseline period in the home cage, sporadic and low-frequency firing patterns predominate, reflecting a familiar and unstimulated environment. Filtered data (300–10,000 Hz) were used for population spike analyses.

**Movie S2. (separate file) Post-experience ripple firing and silent period events in CA1.** This movie displays CA1 multi-unit activity recorded following restraint stress (150–10,000 Hz). The background audio corresponds to spike firing. Frequent, brief ripple firing events alternated with silent periods, illustrating pronounced post-experience modulation of CA1 population activity. Individual ripple firing events appeared visually distinct, consistent with the event-level variability quantified in Fig. S4. Filtered data (300–10,000 Hz) were used for population spike analyses.
